# Four centuries of commercial whaling eroded 11,000 years of population stability in bowhead whales

**DOI:** 10.1101/2024.04.10.588858

**Authors:** Michael V. Westbury, Stuart C Brown, Andrea A. Cabrera, Hernán E Morales, Jilong Ma, Alba Rey-Iglesia, Arthur Dyke, Camilla Hjorth Scharff-Olsen, Michael B. Scott, Øystein Wiig, Lutz Bachmann, Kit M. Kovacs, Christian Lydersen, Steven H. Ferguson, Fernando Racimo, Paul Szpak, Damien A. Fordham, Eline D. Lorenzen

**Affiliations:** Globe Institute, University of Copenhagen, Denmark; The Environment Institute and School of Biological Sciences, University of Adelaide; Center for Macroecology, Evolution and Climate and Center for Mountain Biodiversity, Globe Institute, University of Copenhagen, Copenhagen, 1353, Denmark; Bioinformatics Research Center, Aarhus University; Department of Anthropology, McGill University; Department of Anthropology, Trent University; Natural History Museum, University of Oslo; Norwegian Polar Institute, Fram Center, N-9296, Tromsø, Norway; Fisheries and Oceans Canada, Winnipeg, Canada

## Abstract

The bowhead whale, an Arctic endemic, was heavily overexploited during commercial whaling between the 16th-20th centuries^1^. Current climate warming, with Arctic amplification of average global temperatures, poses a new threat to the species^2^. Assessing the vulnerability of bowhead whales to near-future predictions of climate change remains challenging, due to lacking data on population dynamics prior to commercial whaling and responses to past climatic change. Here, we integrate palaeogenomics and stable isotope (*δ*^13^C and *δ*^15^N) analysis of 201 bowhead whale fossils from the Atlantic Arctic with palaeoclimate and ecological modelling based on 823 radiocarbon dated fossils, 151 of which are new to this study. We find long-term resilience of bowhead whales to Holocene environmental perturbations, with no obvious changes in genetic diversity or population structure, despite large environmental shifts and centuries of whaling by Indigenous peoples prior to commercial harvests. Leveraging our empirical data, we simulated a time-series model to quantify population losses associated with commercial whaling. Our results indicate that commercial exploitation induced population subdivision and losses of genetic diversity that are yet to be fully realised; declines in genetic diversity will continue, even without future population size reductions, compromising the species’ resilience to near-future predictions of Arctic warming.

## Main text

Humans have relied on cetacean species to support their livelihoods for millennia, with whale bones being common at many coastal archaeological sites ^3^. In the Arctic and subarctic, subsistence harvesting of cetaceans started with the arrival of the Thule culture ∼1,000 years ago ^4^, and remains significant to communities across the Arctic. In particular, the bowhead whale (*Balaena mysticetus*) – the only baleen whale found in the Arctic year round – was harvested relatively heavily by the Thule ^5^. However, this harvest may have had little impact because non-breeding whales (mainly yearlings) were predominantly targeted ^6^. Major anthropogenic-driven population declines of bowhead whales likely did not occur before the introduction of commercial-scale harvesting in 1540 CE ^7^.

The bowhead whale was one of the first whale species to be commercially exploited, beginning with whaling by the Basques in the Strait of Belle Isle in southern Labrador, Canada. After depletion of the whales around Labrador, the hunt moved east to Svalbard (Norway) in 1611 ^8,9^ and in 1847-1849, commercial whalers found bowhead whales in the North Pacific, in both the Sea of Okhotsk and the Bering Sea ^1^.

The main reason for commercial exploitation of bowhead whales was the value of the oil rendered from their blubber, which comprises 45-55% of the weight of an individual ^10^. Whale oil was the main source of light in cities across Europe and the eastern United States until the mid 1800s, when gas and later petroleum became available ^1^. By the early 20th century, when commercial bowhead whaling ceased to be profitable, populations had been driven close to extinction ^1^. Bowhead whale protection was put in place in 1931 with the signing of the ‘Convention for the Regulation of Whaling’ ^7^, which banned the harvest of all species in the right whale family (Balaenidae).

Bowhead whales are a major predator of copepods and other zooplankton, and a keystone species responsible for nutrient cycling in Arctic ecosystems. They live in tight association with sea ice, which they rely on for seasonal food resources ^11^ and for protection from killer whales (*Orcinus orca*) ^12^. The decimation of bowhead populations caused by commercial whaling has likely had wide-reaching effects on Arctic marine food webs^13^, and in addition had wide-reaching impacts on Indigenous populations reliant on these ecosystems ^14,15^. Consequently, the future distribution and abundance of bowhead whales is projected to be negatively impacted by on-going declines in sea-ice cover caused by anthropogenic climate change ^16^.

The distribution and genetics of contemporary bowhead populations provides valuable information on current levels of population subdivision and genetic diversity ^17–20^. Furthermore, although genetic data from contemporary bowhead whale populations have been used to study the genetic impacts of whaling ^19,21,22^, demographic inferences may be confounded by the duration of the whaling bottleneck, which was relatively short, considering the long generation time of the species (35-50 years ^17^). Thus, the full genetic consequences of whaling are unlikely to be fully evident using present-day data alone. An accurate representation of long-term, pre-whaling populations is essential to reliably predict the near-future resilience of bowhead whales to a continuously warming Arctic. Only by understanding longer-term patterns of population diversity and subdivision, and by addressing the response of bowhead whales to previous environmental changes, can we assess the degree of genomic erosion associated with commercial whaling, and evaluate the potential of bowhead whales to cope with future change.

Previous studies based on mitochondrial DNA from ancient and present-day bowhead whales have attempted to assess the impact of past climate changes and of commercial whaling on these populations ^23–25^. In these studies, neither climate nor whaling appear to have left a genetic impact. However, mitochondrial DNA represents just a single maternally inherited locus, raising questions as to whether there was no impact, or whether the data analysed lacked sufficient power to detect the genetic effects of these processes.

Bowhead whales have an exceptional – and, in the context of other marine mammals, unprecedented – fossil record. Bowhead whales usually float when dead ^26^, and carcasses end up on shore more frequently than is the case for other large cetaceans that tend to sink to depth when they die. Large numbers of bowhead bones discovered on beaches in the Canadian Arctic Archipelago and in the Svalbard Archipelago (Norway) extend across the past 11,000 years and provide a detailed chronology of bowhead occurrence in these regions. The unprecedented spatial and temporal extent of fossil remains have enabled their use as proxies to estimate the timing and extent of Holocene changes in sea-ice cover in sectors of the Atlantic Arctic ^27,28^.

Using a multi-faceted approach integrating ancient biomolecular analyses (ancient DNA/palaeogenomics, stable isotope analysis) with palaeoclimate data, ecological models, and genomic simulations, we investigated the past 11,000 years of bowhead eco-evolutionary history in the Atlantic Arctic, building on inferences from the exceptional Holocene fossil chronology of > 800 radiocarbon-dated fossil remains, of which 151 dates are new to this study (Figure 1). Specifically, we (i) establish the pre-whaling, long-term demographic trends of bowhead whales, (ii) elucidate their genomic and palaeoecological responses to Holocene climate change events, and (iii) evaluate the long-term genomic impact of commercial whaling.

**Figure 1.**
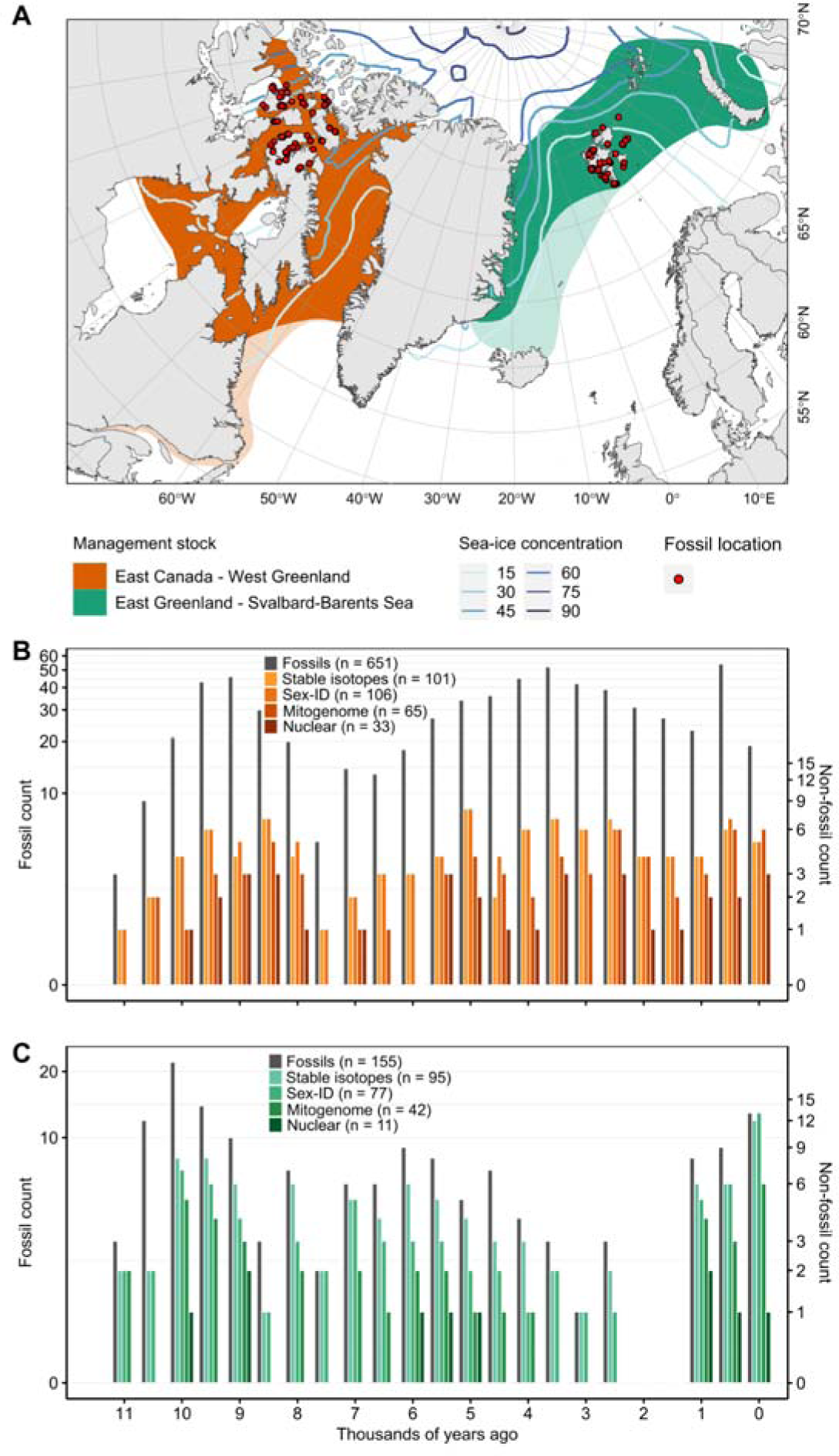
Sample localities and ancient biomolecular data for the Holocene bowhead whale fossil assemblage. (**A**) Map showing the geographic range of the two recognised bowhead management stocks (or breeding populations) in the Atlantic Arctic. Faded colours show the historical distribution that was lost after commercial whaling. Current sea-ice concentrations across the Arctic Ocean are indicated in blue lines. The locality of the Holocene fossil samples from which we retrieved ancient biomolecular data (*δ*^13^C and *δ*^15^N stable isotopes, ancient DNA) are shown as red dots. The complete dataset from (**B**) the Canadian Arctic Archipelago and (**C**) the Svalbard archipelago, including the total number of radiocarbon dated fossils, stable *δ*^13^C and *δ*^15^N isotopes, and ancient DNA for genetic sexing, nuclear genomes (>0.20x coverage), and mitochondrial genomes (>10x coverage) is shown in 500-year time bins.

### Holocene environmental changes and habitat suitability

High-resolution palaeoclimate reconstructions of Holocene environmental change in the Canadian Arctic and Svalbard archipelagos reveal fluctuations in average sea-surface temperature (SST) and sea-ice cover during the past 11,000 years, with the most dramatic changes occurring around Svalbard (Figs 2A,B). According to projections from the HadCM3B-M2.1 coupled general circulation model, the largest shifts in SST and sea-ice cover took place in the early Holocene, with large peaks in SST and troughs in sea-ice cover in the Canadian Arctic Archipelago at ∼10–8.5 thousand years ago (kya). This timing roughly coincides with the onset of the Holocene Thermal Maximum ^29,30^, and the opening of the Nares Strait that connected Baffin Bay and the Lincoln Sea to the Arctic Ocean and flooded the region with nutrient-rich water from the Atlantic some 9 kya ^31^.

**Figure 2.**
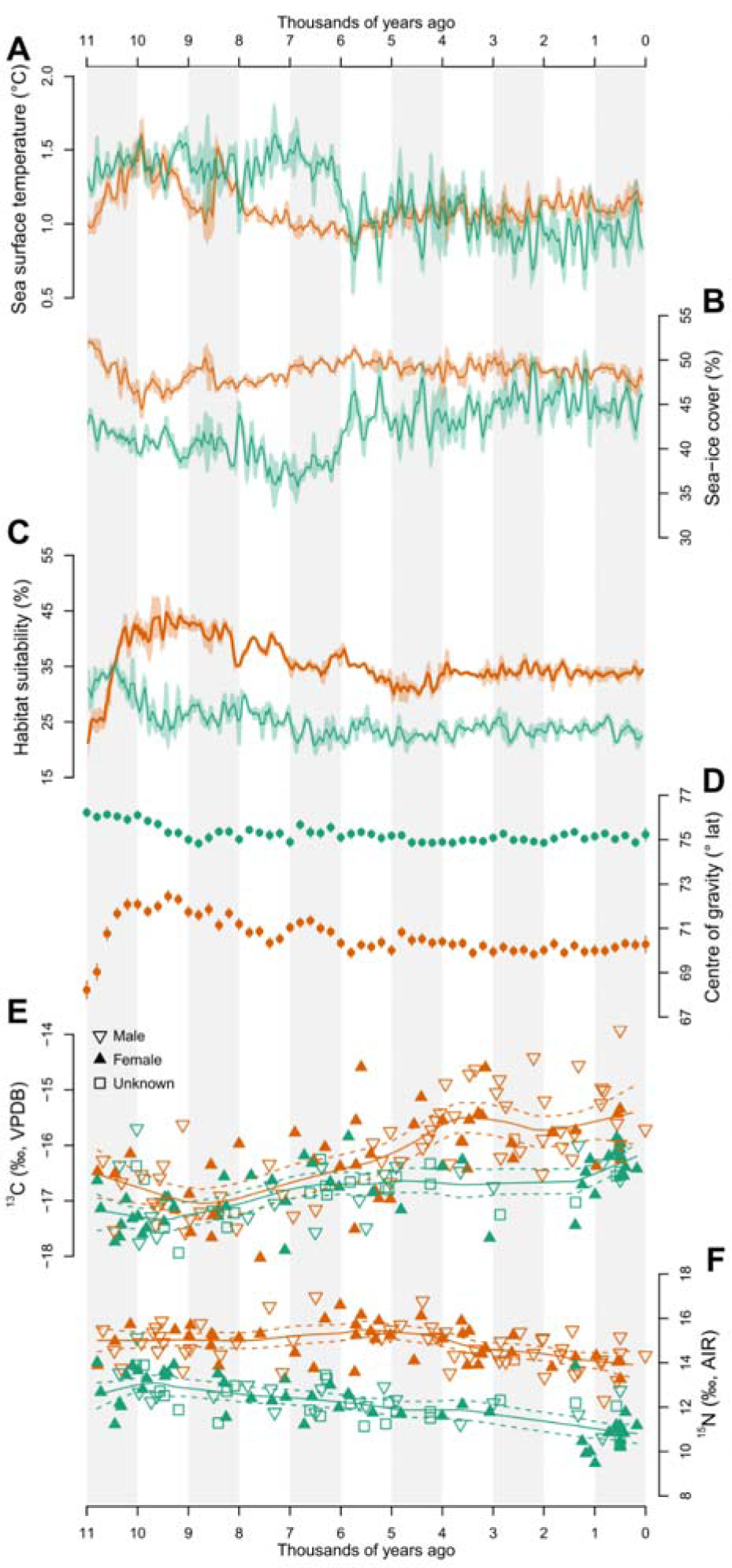
Holocene environmental change, habitat suitability, and palaeoecology. Holocene (**A**) sea-surface temperature (SST) and (**B**) sea-ice cover. (**C**) Percentage of the predefined region containing suitable habitat for bowhead whales based on our models. (**D**) average latitude of suitable habitat within the predefined region. Bone collagen (**E**) *δ*^13^C and (**F**) *δ*^15^N stable isotope values from 196 Holocene bowhead whale fossils, and their genetic sex, if available; the specimens included 94 females, 78 males, and 24 individuals for which sex could not be determined. Trend lines are from a local weighted regression smoothed to fit our scatterplot data. Information about the Canadian Arctic Archipelago is shown in orange and from the Svalbard Archipelago is shown in green.

Differences in the timing of the palaeoclimatic shifts in the western and eastern sectors of the Atlantic Arctic probably reflect heterogeneity in the timing and duration of the Holocene Thermal Maximum across the Arctic ^30^. The rapid decline in SST observed in Svalbard at ∼6 kya may signal the end of the Holocene Thermal Maximum in the region (Fig 2A). However, despite pronounced early Holocene fluctuations in SST and sea-ice cover, our ecological models do not suggest concurrent changes in the estimated area (Fig 2C) or spatial distribution (measured as average latitude, Fig 2D) of suitable bowhead whale habitat. Conversely, habitat suitability projections are relatively stable across time (Supplementary video S1), suggesting long-term ecological resilience of bowhead whales to the climatic perturbations of the Holocene.

In agreement with our spatiotemporal estimates of suitable habitat, we observe limited changes in genetic diversity of bowhead whales across the Holocene, as measured by genome-wide single nucleotide polymorphism (SNP) heterozygosity and nuclear and mitochondrial nucleotide diversity (*π*) assessed in 1,000-year time bins (Fig 3A, Supplementary figs S1, S2, Supplementary tables S1 - S4). This suggests that Holocene environmental shifts, as identified by changes in SST and sea-ice cover, had negligible impacts on the population abundance of bowhead whales. Alternatively, any local changes in genetic diversity were buffered by migration and/or gene flow, which is supported by our genomic simulations (Supplementary figure S3). Nevertheless, the remarkable long-term stability observed in bowhead genetic diversity across the Holocene, despite significant environmental perturbations, is in stark contrast with other Arctic marine mammals for which Holocene demographic reconstructions are available. For example, Greenlandic polar bears (*Ursus maritimus*) experienced marked declines in area of suitable habitat and in population size in the Atlantic Arctic during the Holocene, in response to increasing SST and decreasing sea-ice cover ^32^.

**Figure 3.**
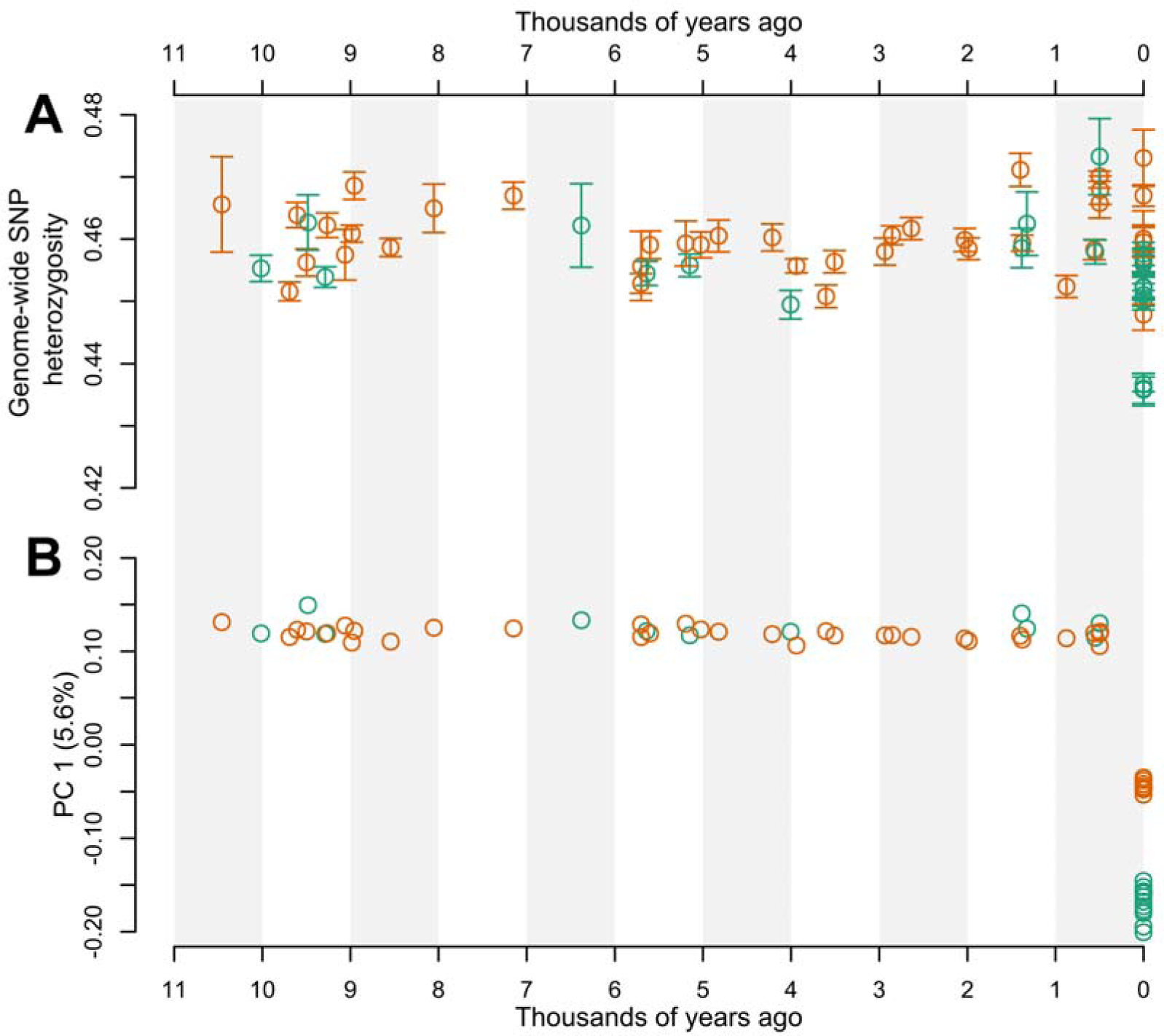
Spatiotemporal patterns of genetic diversity and population subdivision in bowhead whales across 11,000 years. (**A**) ‘Corrected’ individual genome-wide nuclear SNP heterozygosity. Heterozygosity values were corrected in the Holocene fossil individuals by simulating ancient DNA damage patterns onto contemporary individuals and calculating the deviation from the high quality version of the same individual. (**B**) The first axis of a principal component analysis, plotted against the age of each sample. The analyses were based on genome-wide data from 44 Holocene fossil individuals with at least 0.2x coverage and 19 contemporary individuals. Samples from the Canadian Arctic Archipelago are shown in orange (33 pre-whaling Holocene, 7 contemporary), and samples from the Svalbard Archipelago are shown in green (11 pre-whaling Holocene, 12 contemporary).

### Spatiotemporal patterns of bowhead palaeoecology

To further test the response of bowhead whales to environmental change, we analysed bone collagen stable carbon (*δ*^13^C) and nitrogen (*δ*^15^N) isotope compositions from our fossil chronology (Fig 2E,F, Supplementary figure S4). These data provide a proxy for assessing temporal shifts in resources at the base of the food web, i.e., in primary productivity or nutrients, which can be driven by climatic and environmental change ^33^. Bowhead whales are specialised low trophic zooplankton feeders, and thus it is unlikely they would shift trophic position during the time investigated by our study. Sea-ice microalgae, growing within and under sea ice, and phytoplankton, growing in open water, experience regional shifts in community composition and abundance, in response to reductions in sea-ice cover and thickness that alter primary production regimes ^34^. These shifts at the base of the food web drive bottom-up changes in ecosystem structure and function, affecting pelagic secondary production. Such shifts are reflected in the tissue isotopic and chemical compositions of consumers, including bowhead whales ^35^.

Our analysis did not reveal any clear association between spatiotemporal changes in SST or in sea-ice cover and bowhead *δ*^13^C and *δ*^15^N signatures (Fig 2); these results were consistent regardless of the sex of the individual (Supplementary figure S4). Our findings show similar *δ*^13^C values between the western and eastern sectors of the Atlantic Arctic until ∼6 kya, when *δ*^13^C increased in individuals from the Canadian Arctic Archipelago relative to the Svalbard Archipelago. *δ*^13^C values in specimens from the Canadian Arctic Archipelago continued to rise until ∼3.5 kya. The timing of the onset of the increase corresponds with the end of the Holocene Thermal Maximum, which may have caused a shift in primary producers as temperatures changed. A similar increase in *δ*^13^C has also been documented in Northwest Greenland in sedimentary organic carbon ^36^, suggesting this pattern is not specific to bowhead whales, but rather reflects a regional shift in primary productivity in the western sectors of the Atlantic Arctic.

We carried out a genomic analysis of nuclear SNPs to find those with the highest likelihood of change in allele frequency associated with time. This revealed 18 sites located within 16 annotated genes, linked to an array of potential phenotypes that may be associated with responses to environmental changes in the Arctic, including body size, metabolism, regulation of cardiovascular and renal systems, and development of adipose tissue (Supplementary tables S5, S6). Visualising changes in allele frequency through time reveals an allelic shift at most of the sites at ∼5-3 kya (Supplementary figure S5). The relative conformity in timing of allele changes, despite the sites being present in vastly different parts of the genome, suggests a large change in selective pressure across the species range. The timing coincides with the divergence in the trajectory of *δ*^13^C values between the two regions, and hence supports an ecological driver of change (Fig 2E).

The clear differentiation in *δ*^15^N between the Canadian Arctic Archipelago and the Svalbard Archipelago is likely due to regional variability in *δ*^15^N at the base of the food web, as has been reported in other marine predators ^37–39^. During the second half of the Holocene, *δ*^15^N in bowhead whales around both the Canadian Arctic Archipelago and the Svalbard

Archipelago decreased gradually (Fig 2F), suggesting a slow change in nutrient dynamics, possibly decreasing rates of sedimentary denitrification, a process which results in ^15^N-enrichment in water column organic matter in Arctic and subArctic continental shelf environments ^40^. Such a change would be reflected in the *δ*^15^N of consumers, such as bowhead whales ^41^.

### Spatiotemporal patterns of genomic structuring

Bowhead populations are recognised by the International Whaling Commission (IWC) as comprising four geographically segregated stocks, based on genetics and non-genetic data, including telemetry ^42^. Contemporary bowhead whales in the Canadian Arctic Archipelago belong to the ‘East Canada West Greenland’ stock (Fig 1A). Contemporary bowhead whales from around the Svalbard Archipelago belong to the ‘East Greenland Svalbard Barents Sea’ stock. Our *F*_ST_ analysis shows the level of genetic differentiation between the two stocks is ∼3.7x higher at present than during the Holocene, indicating that population subdivision between stocks is a recent phenomenon (Supplementary figure S6). Indeed, we found no indications of genomic population subdivision in our fossil individuals, suggesting bowhead whales comprised a single panmictic population during the Holocene (Fig 3B and Supplementary figures S7 - S12). However, our observed regional differences in *δ*^15^N indicate whales mostly fed locally, in the western or eastern sector of the Atlantic Arctic, where their fossils were found (Fig 2F), suggesting geographic segregation of individuals in both archipelagos. These divergent findings may be reconciled if sufficient levels of connectivity across the Atlantic Arctic were maintained throughout the Holocene, to facilitate gene flow and prevent genetic subdivision. This scenario is supported by genomic simulations, which show no genetic subdivision when the populations are connected by as few as ∼5 migrants per generation (Supplementary figure S3).

Our ecological modelling showed a high likelihood of suitable habitat around northern and southern Greenland, connecting the Canadian Arctic Archipelago and the Svalbard Archipelago from 11,000 years ago until 1950 (the most recent time point in our models) (Supplementary video S1). Movement of bowhead whales between the western and eastern sections of the Atlantic Arctic north of Greenland is supported by Holocene fossil evidence (Supplementary video S1). Although there is no fossil evidence from the southern coast of Greenland to support this, the absence of fossil remains could reflect the acidity of substrates in the region, which likely limits the preservation of organic material, potentially masking whale presence.

### The genomic impacts of commercial whaling

At the onset of commercial whaling, the range of bowhead whales extended further south of their current winter range in northern Labrador, Canada (Fig 1A) ^43^. Their range included the Strait of Belle Isle and the Gulf of St Lawrence, which were the first areas where bowhead whales were heavily hunted by Basque whalers ^1^. This extension of their historical range so far south of their current distribution could be due to their relative displacement during the cooler climates of the Little Ice Age (∼1300-1900 AD). The disruption of gene flow between contemporary stocks relative to their pre-whaling counterparts (Fig. 3B, Supplementary figure S6) could in part be explained by the extirpation of bowhead whales from the southern parts of their range by early commercial harvests ^1^.

During the four centuries of commercial bowhead whaling, the cumulative offtakes in the eastern sector of the Atlantic Arctic are estimated to have been much more severe than in the western sector ^1^. The ‘East Greenland Svalbard Barents Sea’ stock numbered in excess of 52,500 bowhead individuals prior to whaling ^9^. Contemporary bowhead whales in the region number some few hundred individuals ^9,44^, suggesting a >98% population size decline. In contrast, the pre-whaling estimates for the ‘East Canada West Greenland’ stock is ∼18,500 individuals ^45^, and the current population numbers ∼6,000 individuals ^46^, equivalent to ∼70% population size decline. The difference in whaling intensity between regions is mirrored in our dataset; contemporary ‘East Greenland Svalbard Barents Sea’ individuals have significantly lower mean genome-wide SNP heterozygosity and genome-wide nucleotide diversity compared to Holocene individuals (our analysis suggests ∼2% loss for each estimate; Fig 3A, Supplementary figure S1, Supplementary tables S1, S3). In contrast, contemporary bowhead whales from Canada have diversity values that do not significantly differ from Holocene individuals from the Canadian Arctic Archipelago, indicating they reflect long-term pre-whaling diversity more closely than their Svalbard counterparts.

The genetic impact of whaling is also visible in other parameters in the genomes of the ‘East Greenland Svalbard Barents Sea’ stock. Our demographic reconstruction based on 12 contemporary samples from the Svalbard Archipelago estimated ∼12% loss in effective population size (Ne) ∼5 generations ago, which is equivalent to 175-250 years ago (with a generation time of 35-50 years ^17^; Supplementary figure S13). Similar findings were recently reported in bowhead whales in the ‘East Canada West Greenland’ stock, which show a large decrease in Ne ∼4 generations ago ^20^. However, the reliability of bowhead demographic reconstruction based exclusively on contemporary data is questionable, as inferences of population changes are indirect, and may be confounded by long generation time, short bottleneck duration, and the magnitude of population depletion. Thus, long-term pre-whaling data is imperative for reliably quantifying the relative and long-term genetic impact of commercial whaling.

To estimate the most probable level of Holocene migration between the western and eastern sectors of the Atlantic Arctic, and the magnitude of the bottleneck caused by commercial whaling, we compared our empirical estimates of spatiotemporal changes in genetic diversity and *F*_ST_ (Fig 3A, Supplementary figures S1 and S6) with genomic simulations of various demographic scenarios. Based on a model of our pre-whaling Canadian bowhead population, we estimate that stable genetic diversity would have been maintained if populations were reduced by a maximum of 50% (Fig 4A). Similarly, using a model of a 2% reduction in genetic diversity in a population modelled after our pre-whaling Svalbard bowhead population, we estimate a population decline of 92-96%. To emulate the relative temporal change in *F*_ST_ observed between Holocene and contemporary individuals, our simulations require migration to have been reduced to ∼1 individual per generation, or to have ceased entirely (Fig 4). Thus, our findings suggest the sustained migration levels that buffered changes in genetic diversity during the Holocene (at least ∼5 individuals per generation) are likely no longer happening in contemporary populations. Our spatiotemporal estimates of habitat suitability do not suggest a loss of habitat connectivity south or north of Greenland since the onset of commercial whaling, indicating any cessation or reduction in gene flow is driven by demographic changes rather than environmental changes.

**Figure 4.**
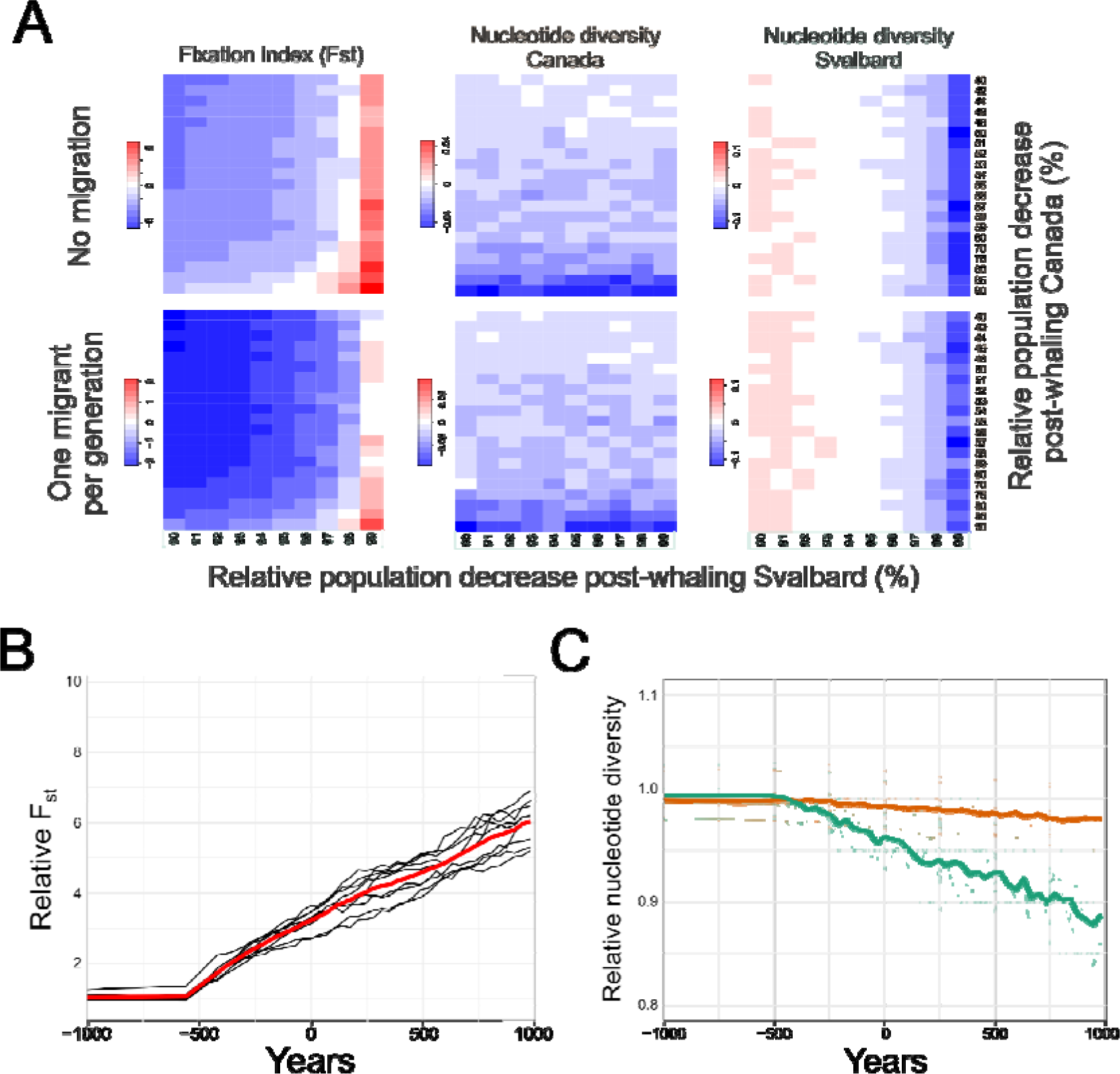
Simulated past and future changes in migration and genetic diversity. (A) comparison of delta (difference between pre- and post-bottleneck) statistics between empirical and simulated data; 0 (in white) indicates simulated parameter combinations that match the mean observed values of the empirical genomic data. Future projections of (B) changes in *F*_ST_ between the two populations and (C) changes in nucleotide diversity within the Canadian Arctic Archipelago (orange) and Svalbard Archipelago (green) based on the simulated data, where the population representing Canada decreased 48% and the population representing Svalbard decreased 97%, and migration between the populations ceased after the bottleneck. Bold lines represent the mean values. All simulations assumed the bottleneck occurred ∼500 years ago and that population size did not change subsequently. Simulations were performed in generations and converted to years, assuming a generation time of 35 years.

Using forward simulations, we show that population divergence and loss in genetic diversity associated with commercial whaling is not yet fully realised in contemporary populations (Fig 4B,C), likely due to the long generation time of bowhead whales and the relatively recent timing of the bottleneck. The diversity of contemporary bowhead whales underestimates the actual loss due to whaling, which ceased in both regions < ∼5 generations ago ^1^, and thus it may take several more generations for the genetic signs of low population size to become evident. If all other demographic factors stay the same, genetic diversity will continue to decrease in both populations, resulting in ∼1% decrease in the ‘East Canada West Greenland’ stock and ∼4% decrease in the ‘East Greenland Svalbard Barents Sea’ stock over the next 100 years or ∼3% decrease in the ‘East Canada West Greenland’ stock and ∼15% decrease in the ‘East Greenland Svalbard Barents Sea’ stock over the next 1,000 years. This is expected, as genetic diversity loss has a time-lag relative to demographic decline, particularly in long-lived species ^47^. Genetic diversity loss in collapsed populations (e.g., Svalbard) is expected to continue even if there is demographic recovery ^48,49^, because small populations continue to pay a ‘genetic drift debt’ ^50^. This sustained genomic erosion, especially in the ‘East Greenland Svalbard Barents Sea’ stock, brings into question the long-term resilience of the population. The loss of genetic diversity will likely be exacerbated by further declines in population size, which are predicted based on modelled future patterns of habitat suitability, which shows a significant decrease and northwards shift towards year 2100 ^16^. Overall, this highlights the pressing need for long-term protection and (genetic) monitoring of bowhead whale populations.

Prior to commercial whaling, bowhead whales in the western sector of the Atlantic Arctic endured hundreds of years of Palaeoinuit subsistence hunting. Thule harvests in the central and eastern Canadian Arctic are difficult to estimate, but have been approximated at ∼11,500 whales between 1200–1529 AD ^5^, roughly 20% of total commercial harvests. The average offtakes translate to ∼40 individuals per year for Thule subsistence harvests, and ∼150 individuals per year for commercial harvests. However, there was a relatively short, concentrated period of significantly higher commercial offtakes in eastern Canada and West Greenland of up to 1500 individuals per year between ∼1830-1840 ^5^.

Based on the size of bones retrieved from archaeological sites, Thule are inferred to have focused almost exclusively on non-breeding individuals (yearlings and small juveniles) ^6^, and thus likely had lower impacts on the species relative to commercial whaling, which was either non-selective, or targeted the largest animals ^51^. Indeed, a negligible impact is supported by our findings, which show no evidence of genetic diversity loss prior to commercial whaling (Fig 3). The longer period of sustained bowhead harvests in the Canadian Arctic Archipelago by Thule and commercial whalers, relative to Svalbard, where there have been no subsistence harvests, makes the regional patterns of diversity loss in contemporary individuals more profound; our data show commercial exploitation left the Svalbard whales in a much worse state than their Canadian counterparts, in agreement with regional differences in offtakes estimates^1^.

The Arctic is experiencing transformative change due to climate warming. We are already observing a four-fold amplification of the rate of change in temperature in this region, compared to the global average ^2^. Climate models predict further increases in SST and losses of sea-ice cover in our two study regions in the near future^16^ (Supplementary tables S7 and S8). While some climatic perturbations that followed the last ice age were similar in pace and magnitude to what is predicted for the 21st century ^52^, absolute temperatures in the Arctic this century are predicted to exceed those experienced during the past 55 million years ^53,54^. We show that bowhead whales across the Atlantic Arctic were resilient to the past 11,000 years of environmental change, with higher levels of nuclear genetic diversity and higher potential for gene flow in the Holocene, perhaps providing the capacity for the species to deal with past environmental perturbations. However, our study indicates commercial whaling eroded genetic diversity, and that we have yet to see the full genomic consequences of the commercial decimation of bowhead populations, which may impact the species’ resilience to near-future climate change.

## Methods

### Bowhead samples

All specimens analysed were previously identified in the field as bowhead whales. For ecological modelling, we compiled a record of available radiocarbon dated subfossil bowhead whales from across the circumpolar Arctic, totalling 824 individuals after filtering ^23,27,55–84^ (Supplementary table S9). We included only fossils that were known to be from bowhead whales, and excluded samples processed before 1980 if we did not have information on whether dating of the specimen was performed using collagen and with adequate pretreatment (especially cleaning), as these protocols were not in regular use until the early 1980s ^85,86^. The largest number of specimens suitable for ecological modelling were from the Canadian Arctic Archipelago and Svalbard Archipelago chronologies; a subset of these samples were analysed using ancient biomolecules, mentioned above and detailed later. We included 71 new radiocarbon dates of bowhead fossils from around the Svalbard archipelago, and East Greenland (Supplementary table S9). The samples were identified and dated following Wiig et al. 2019 ^55^. We also included 80 new radiocarbon dates from bowhead whale fossils from around the Central Canadian Archipelago, which were identified and dated following Dyke et al 1996 ^27^.

To ensure comparability between sample ages across our analyses, we recalibrated all original radiocarbon dates with Calib v7.0.4 using the marine13 calibration curve, a specified age span of 100 years, and unique marine reservoir correction (delta R) values depending on the region in which the specimen was found. For samples from the Canadian Arctic (n = 652) we used a delta R of 170±95 ^87^; from around Alaska (n = 4), we used a delta R of 506±83 ^88^; from the Svalbard Archipelago and the Norwegian coast (n= 167), we used a delta R of 7±39^89^.

For ancient DNA and stable isotope analyses, we sub-sampled radiocarbon dated bowhead whale bone specimens from the Canadian Museum of Nature, Ottawa (Canadian Arctic Archipelago samples) and Natural History Museum, University of Oslo (Svalbard Archipelago samples). All except one Canadian sample (>33 kya) and three Late Pleistocene Svalbard samples were dated to the Holocene. An overview of the samples and their associated biomolecular data is included in Supplementary tables S10 (stable *δ*^13^C and *δ*^15^N isotopes), and S11+S12 (ancient DNA).

Contemporary bowhead whales are recognised as belonging to four distinct management units (termed stocks or breeding populations), based on genetics and non-genetic data (incl. telemetry) ^42^. To contextualise the Holocene ancient DNA data, we included comparable genomic data from samples collected from the two contemporary stocks in the Atlantic Arctic. We generated genomic data from seven samples from the contemporary ‘East Canada West Greenland’ stock, and downloaded published genomic data from 12 bowhead individuals from the ‘East Greenland Svalbard Barents Sea’ stock (sampled 2017-2018, Genbank bioproject: PRJNA643010) ^17^. Sample and data overviews for the contemporary samples are provided in Supplementary table S13.

### Ecological niche modelling

#### Climate data

Paleoclimate data were accessed using a high resolution (1° x 1°) oceanic climate dataset for the period 60 thousand years ago (kya) through to the present (1950 C.E.) ^90^. These data were generated by temporally linking discrete snapshot simulations from the HadCM3B-M2.1 coupled general circulation model ^91^. The HadCM3B-M2.1 model has a nominal oceanic resolution of 1.25° × 1.25° and is run as a series of snapshots at 1000-year intervals between 0 (1950 C.E.) and 22,000 BP, and 2000-year intervals between 22,000 BP and 60,000 BP. The snapshot simulations have been linked using splines based on monthly climatologies, before interannual and millennial scale variability (e.g. Dansgaard-Oeschger ^92^ and Heinrich ^93^ events) was imposed on the timeseries. The data has been downscaled to the final 1° × 1° resolution using bilinear interpolation. Tests of the HadCM3B-M2.1 model show that it reproduces global and regional sea-surface temperatures and surface salinity ^91^. While no validation of the HadCM3B-M2.1 sea-ice dynamics has been done, a validation using the same underlying model (HadCM3), has shown a good fit to observed sea-ice extents and declines ^94^.

Simulation of the climate system by even the most advanced global climate models contain notable biases ^95^. Consequently, it is crucial to address these model biases in order to achieve realistic paleoclimate simulations for use in studies of long-term ecological dynamics ^96^. Our climate data were bias-corrected using an additive delta (change-factor) method ^95^ for sea-surface temperature, and a multiplicative correction for sea-surface salinity and sea-ice concentration. Sea-surface temperature and salinity were bias corrected against the World Ocean Atlas 2018 dataset (https://www.ncei.noaa.gov/products/world-ocean-atlas), with sea-ice concentration corrected against the Twentieth Century Reanalysis dataset ^97^ using a climatological period of 1850-1950 C.E. Multiplicative bias corrections were capped at 3x the simulated value ^90^. Corrected sea-ice cover values that exceeded 100% were truncated back to 100%. No bias-correction was done on sea-ice thickness due to there not being a suitable dataset covering the end of the model simulation period (1850-1950 C.E.).

The resulting paleoclimate simulations were a continuous time series of maps of climatological monthly averages, calculated over a 30-yr window, with a step of 50 years, for the period 60 kya to 0 kya. We extracted seasonal data for sea-surface temperature (SST; °C), sea-surface salinity (SAL, ‰), sea-ice cover (SIC, %), and sea-ice thickness (SIT, m) for our study region. This data was then used to generate 30-yr averages of four variables (for each season) at 50 year timesteps: (i) seasonal mean SST; (ii) seasonal mean SAL; (iii) seasonal mean SIC; and (iv) seasonal mean SIT. All fossil records for bowhead whales were then matched to this data and used to calibrate an ecological niche model (see below).

Following exploratory data analyses we opted to use summer (June, July, August) SST, SAL, SIC in our ecological niche model. We did not use summer SIT as we were unable to bias-correct it. These metrics are theorised to be as successful as more direct (proximal) variables in predicting the relationships between environmental pattern and process, particularly in extreme environments where species are not occupying optimal parts of their potential realised niches ^98–100^. The choice of summer seasonal data is justified as we have a large number of samples above the northern edge of the contemporary Canadian Arctic Archipelago population boundary (Fig 1), and bowhead whales from this region are known to move northwards in summer following the sea ice as it retreats ^27^. Furthermore, bowhead whales in the Svalbard area have been observed to be on average ∼100 km offshore during the summer ^101^. Given that we only used radiocarbon dated fossil records - primarily located along coastlines - to calibrate and validate our ecological models, Svalbard fossils in all likelihood resulted from animals that died in summer.

Previous work has shown that bowhead whales prefer cold, ice-covered water, with individuals spending most of their time in a narrow temperature range > -1°C SST < 1°C, near the marginal ice zone, but also moving into areas with >90% SIC during winter ^101^. Proximity to coastline, and consequently, bathymetric depth has also been shown to be an important variable controlling bowhead whale distribution ^101^, but we were not able to calculate these metrics (e.g. bathymetric depth, distance from continental shelf) accurately because accurate bathymetric data for our high-resolution paleoclimate reconstructions do not exist ^90^. We included salinity as a proxy for regional differences in ocean productivity, with decreased pelagic and benthic diversity often occuring in areas of lower salinity ^102^. Consequently, our estimates of SST, SAL, and SIC metrics could be considered both proximal and distal predictors as they have a direct (proximal) influence on bowhead whale physiology and behaviour (and therefore fitness), and an indirect (distal) influence on prey distributions.

#### Ecological niche model

We created an ecological niche model (ENM) for bowhead whales using the Hypervolume package for R ^103^. We generated best estimates of the ecological niche as a 3-dimensional hypervolume ^104^ across time ^99^. Hypervolumes were constructed using the “Gaussian” hypervolume method ^105^, with bandwidths, number of standard deviations, and the probability threshold tuned using independent calibration and validation datasets. Gaussian hypervolumes were built by defining a Gaussian kernel density estimate on an adaptive grid of random n-dimensional points around the original data points. The bandwidth multiplier, number of standard deviations, and the probability threshold all control the size and configuration of the kernel density estimate ^103,105^. We withheld a stratified 10% of our expanded occurrence records to use as an independent validation set, with the remaining 90% of records used to calibrate the hypervolume. We intersected the 30-year averages for our climate and environmental variables for each georeferenced fossil for the period ± 2 SD around the estimated age of the fossil, ensuring that each fossil record had a time series of climate data associated with it ^106^. This time series represents the period over which bowhead whales were likely to have been present near the fossil sites, given inherent dating uncertainty. Before pairing the fossil records with the environmental and climate data (see above) to define the niche, we merged records where there was spatiotemporal overlap within each 1° × 1° grid-cell. To do this, longitude and latitude values for fossils (Supplementary table S9) were rounded to one decimal place (retaining ∼11.1 km of accuracy) and grouped. Each record was then checked for temporal overlap with all other records in the same group. Temporal overlap was defined as overlapping confidence intervals for the calibrated radiocarbon ages (Calibrated Age ± 2 S.D.). Where temporal overlap occurred, the confidence intervals were merged for all overlapping records resulting in a single record with an expanded age interval. Pre-processing the collated fossil records using this approach reduced the number of records for modelling the niche to 585 (n = 526 calibration, n = 59 validation). Expanding the calibration and validation datasets to their full temporal coverage resulted in 10,798 calibration records and 1,148 validation records.

To characterise the environmental conditions at each fossil location, SST, SIC, and SAL were calculated as the average values from the nearest ocean grid-cell containing the fossil and the 8-nearest cells. The 9-cell averaging approach was chosen to minimise fine-scale artificial accuracy/biases introduced during the bias correction and downscaling of the climate data ^90^ and to overcome positional uncertainty regarding the potential ocean cells from which the fossils were likely to have arisen. For this process, fossils that were located on land according to the temporally explicit land/sea mask, were snapped to their nearest ocean-cell, up to a maximum distance of 150 km, before the nearest 8-cells to the “new” fossil location were identified (Supplementary figure S14).

#### Climate suitability projections

Spatially and temporally explicit projections of habitat suitability were created at 50-year generational time steps from 11 kya BP to 0 BP for bowhead whales. We opted to set an upper limit on our hindcasts as only 7% of our fossil record was from fossils older than 11 kya BP and we therefore had reduced confidence in projections of habitat suitability before this time. Comparisons between spatial projections of habitat suitability from the hypervolume package, and more common maximum entropy methods ^107^ have shown similar results ^105^.

Using the full multi-temporal fundamental niche hypervolume, we used the 10% validation test set to fine-tune the two parameters that affect the probability density (i.e. habitat suitability) of the hypervolume: (i) weight.exponent, and (ii) edges.zero.distance.factor. These two parameters in combination control the rate and distance at which habitat suitability shifts to 0 from its empirical maximum. A grid search was done using both parameters (edges.zero.distance.factor range = 1:10; weight.exponent range = -1:-3), before extracting values of habitat suitability at fossil locations (temporally explicit), and then calculating the Boyce Index and area-under the receiver operating curve (AUC) values ^108,109^. The Boyce index is a presence-only evaluation measure used to discriminate how much projections of habitat suitability at presence locations differ from random expectation, with higher Boyce values indicating greater habitat suitability at presence locations than would be expected by chance. Likewise, higher AUC values indicate greater capacity for the hypervolume projections to discriminate between background locations and training/validation sites. Background points for calculating both measures were defined using the background points from the full hypervolume. Final parameters were chosen based on the combination of parameters that maximised the Boyce Index and AUC and consequently habitat suitability at fossil locations through time. Following tuning and validation, the weight.exponent was set to -1 and edges.zero.distance.factor was set to 3. As the outputs of the hypervolume_project function are not bound by [0, 1]^103,105^, each of the projections of habitat suitability was then rescaled to the range 0-1 using the 10% training presence threshold (OR10). The OR10 is defined as the threshold which excludes regions with suitability values lower than the values for 10% of training records – the assumption being that the lowest 10% of training records are from regions that are not representative of the species overall habitat and can be omitted. Values were rescaled using the formula:

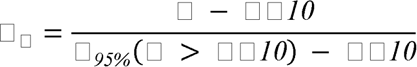

Where X_r_ is the rescaled suitability value, x is the original value, OR10 is the 10^th^ percentile omission threshold from the training data, and P_95%_ is the 95^th^ percentile of all x values > OR10. We opted to rescale based on the 95^th^ percentile of suitability values due to the extreme right skew of the suitability values inflating the maximum value. Projections of habitat suitability were then reprojected using bilinear interpolation to a stereographic polar projection with a resolution of 100 km x 100 km.

### Stable isotope analysis

Approximately 100 mg of powdered bone was removed from subfossil bowhead whale specimens and demineralized in 0.5 M HCl for 4 h under constant motion (orbital shaker). The samples were then rinsed with Type I water (18.2 MΩ·cm), treated with 0.1 M NaOH for 20 minutes. This step was repeated until there was no colour change in the solution. The samples were then heated at 75°C for 36 hrs in 3.5 mL 0.01 M HCl to solubilize the collagen, and freeze-dried.

We determined carbon and nitrogen isotopic and elemental compositions using a Nu Horizon isotope ratio mass spectrometer (IRMS) coupled to a EuroVector 3000 elemental analyzer (EA). The *δ*^13^C and *δ*^15^N values were calibrated relative to the international reference scales (VPDB and AIR) using USGS40 and USGS41a ^110,111^. We assessed measurement uncertainty using three in-house standards with the following established isotopic compositions: SRM-1 (caribou bone collagen, *δ*^13^C = *□*19.36±0.11 ‰, *δ*^15^N = +1.81±0.11 ‰), SRM-2 (walrus bone collagen, *δ*^13^C = *□*14.77±0.11 ‰, *δ*^15^N = +15.59±0.11 ‰), SRM-14 (polar bear bone collagen, *δ*^13^C = *□*13.67±0.07 ‰, *δ*^15^N = +21.60±0.15 ‰), and SRM-15 (phenylalanine, *δ*^13^C = *□*12.44±0.04 ‰, *δ*^15^N = +3.08±0.12 ‰). To check for homogeneity of the collagen, twenty percent of the samples were analysed in duplicate. Standard uncertainty was calculated to be ±0.14 ‰ for *δ*^13^C and ±0.28 ‰ for *δ*^15^N ^112^.

We generated locally estimated scatterplot smoothing (LOESS) trendlines using the statistical software package PAST v4.03 ^113^ with a smoothing factor of 0.25.

## Genomics

### Ancient DNA data generation

We extracted DNA from our subfossil bowhead whale specimens using a modified version of a previously published protocol ^114^. Modifications included using a modified version of the Qiagen PB binding buffer ^115^ and concentrating the extraction supernatant to ∼100ul using Amicon spin columns prior to purification. We measured the DNA concentration in the extracts using the Qubit high sensitivity kit. We performed a USER enzyme treatment step to remove uracil residues from damaged DNA and the resultant abasic sites ^116^. We built Illumina sequencing libraries from the USER treated extracted DNA following the BEST protocol ^117^, with a predetermined Illumina adapter mix concentration (1 - 50 uM) based on the DNA extract concentration and a set number of indexing PCR cycles, predetermined a priori through a qPCR reaction. Index PCR was performed using dual-indexing and libraries were combined into pools of ∼50 unique indices. Index reactions were performed using the Kapa Hifi Uracil + Readymix and the following PCR conditions: 98°C for 45 seconds, then 98°C for 15 seconds, 60°C for 30 seconds, and 72°C for 20 seconds for the number of cycles predetermined via qPCR, and finally a cool down to 10°C. We sequenced each library pool on a single Illumina Hiseq 4000 lane at the GeoGenetics Sequencing Core, University of Copenhagen using 80 bp single end (SE) chemistry. We selected individuals for deeper sequencing based on endogenous DNA content (number of unique mapped reads/total number of raw reads), age, and locality. We built new sequencing libraries for the selected samples which were sequenced on an Illumina Hiseq 400 with 80 bp SE chemistry.

We enriched 34 Svalbard individuals for mitochondrial genomes using RNA baits based on the published bowhead whale mitochondrial genome (KY026773.1). Enrichment was performed using the hybridization capture myBaits Custom DNA-Seq kit (Arbor Biosciences). Following myBaits recommendations, we used between 150 – 280 ng of starting material of each indexed library for every capture reaction. The capture procedure was carried out as described in the myBaits manual v.5.00; we used the High Sensitivity conditions, which are optimised for ancient samples, with a hybridization step at 55 °C for 24 h. Post-capture, the libraries were re-amplified using Kapa Hifi Uracil + Readymix and the following PCR conditions: 98 °C for 45 minutes, then 98 °C for 20 seconds, 60 °C for 30 seconds, and 72 °C for 45 seconds for 14 cycles, and a final elongation at 72 °C. Re-amplified libraries were quantified and quality checked as described above. Sequencing was carried out on a NovaSeq 6000 at Novogene Europe with 150 bp PE chemistry.

### Contemporary genomic data generation

We extracted DNA from the seven contemporary Canadian individuals using a DNeasy blood and tissue kit (Qiagen) following the manufacturer’s protocol. We fragmented the extracted DNA to an average length of ∼450 bp using a M220 Focused-Ultrasonicator™ (Covaris). We built Illumina sequencing libraries from the fragmented extracts using the BEST protocol ^117^, with an Illumina adapter mix concentration of 20uM, and 15 cycles during the indexing PCR step. We cleaned the indexed libraries using a SPRI bead DNA purification method. Each indexed library was sent to Novogene for 10 Gb of 150 bp paired end (PE) sequencing on a Novaseq Illumina platform.

### Data processing

For the 202 subfossil individuals that successfully produced sequencing data, we trimmed adapter sequences and removed reads shorter than 30 bp from the raw reads using skewer v0.2.2 ^118^. We mapped the trimmed reads to the bowhead whale reference genome ^119^ including the mitochondrial genome (Genbank accession: KY026773.1) using Burrows-wheeler-aligner (BWA) v0.7.15 ^120^ utilising the aln algorithm, with the seed disabled (-l 999) (otherwise default parameters). We parsed the alignment files and removed duplicates and reads of mapping quality score <30 using SAMtools v1.6 ^121^. We checked for ancient DNA damage patterns using mapdamage2 ^122^.

For the 19 contemporary individuals (Canada n = 7, Svalbard n = 12), we trimmed adapter and poly-G sequences and removed reads shorter than 30 bp from the raw reads using Fastp v0.20.1 ^123^. We merged overlapping paired-end reads using FLASHv1.2v11 ^124^, using default parameters. We mapped both merged and unmerged reads to the bowhead whale reference genome using BWA with the mem algorithm (otherwise default parameters). We parsed the alignment files and removed duplicates and reads of mapping quality score <30 using SAMtools.

### Nuclear genomes

We found putative sex chromosome scaffolds in the bowhead whale reference genomes by aligning it to the Cow X (Genbank accession: CM008168.2) and Human Y (Genbank accession: NC_000024.10) chromosomes. We performed the alignments using satsuma synteny v2.1 ^125^ with default parameters.

### Relatedness

We assessed whether any of our contemporary individuals could be closely related to each other using NGSrelate v2 ^126^. As input for this we calculated genotype likelihoods for the contemporary individuals using ANGSD v0.921^127^. We calculated genotype likelihoods using the GATK algorithm (-GL 2), specified the output as a binary beagle file (-doGlf 3), and applied the following filters: only include reads with a mapping quality greater than 20 (- minmapQ 20), only include bases with base quality greater than 20 (-minQ 20), only include reads that map to one location uniquely (-uniqueonly 1), a minimum minor allele frequency of 0.05 or greater (-minmaf 0.05), only call a SNP if the p-value is less than 1e^-6^ (-SNP_pval 1e-6), infer major and minor alleles from genotype likelihoods (-doMajorMinor 1), skip triallelic sites (-skipTriallelic 1), remove sex scaffolds and scaffolds shorter than 100 kb (-rf), and call allele frequencies based on a fixed major and an unknown minor allele (-doMaf 2). We determined a relatedness coefficient (RAB) >0.125 (equivalent of first cousins) as closely related.

### Sex determination

We calculated the average coverage of scaffolds aligning to the X chromosome and the autosomes (scaffolds not aligning to either the X or the Y chromosome) using SAMtools depth. We determined the sex of an individual by calculating the X:A, the ratio of coverage on the X scaffolds to the autosomal scaffolds. Of the 202 individuals analysed, 16 subfossil individuals had <5,000 mapped reads and were not considered further for sex determination analysis^128^. If an individual had an X:A ratio of <0.7 it was designated as a male. If an individual had an X:A ratio of >0.8 it was designated as a female. Individuals with ratios between 0.7 and 0.8 were deemed undetermined^128^. To investigate changes through time, we subsequently pooled individuals into 1,000 year time bins and calculated the ratio of males to females.

### Population structure

We investigated population structure by performing Principal Component Analyses (PCA) using PCAngsd v0.95 ^129^ using all individuals with >0.2x genome-wide coverage. As the input for PCAngsd, we generated a genotype likelihood beagle file in ANGSD using the following parameters: -minmapQ 30, -minQ 30, -GL 2, -doGlf 2, -doMajorMinor 1, remove transitions (-rmtrans 1), -doMaf 2, -SNP_pval 1e-6, -minmaf 0.1 -skiptriallelic 1, -uniqueonly 1, only include sites where at least 40 individuals have coverage (-minind 40), only including autosome scaffolds <100kb (-rf).

We tested the robustness of our PCA results to coverage and aDNA damage patterns by repeating the analyses with modified versions of the 12 high-coverage contemporary ‘East Greenland Svalbard Barents Sea’ individuals, while keeping all factors, including parameters, unmodified. To test the impact of coverage, we downsampled all 12 contemporary ‘East Greenland Svalbard Barents Sea’ individuals to 2x using SAMtools. To test the impact of shorter read lengths and aDNA damage patterns, we trimmed the forward reads of the 12 individuals to 80 bp using skewer, and simulated aDNA damage patterns on the ends of the reads, using TAPASv1.2 ^130^, based on the mean damage misincorporation values generated by Mapdamagev2 from all ancient bowhead samples. We mapped the simulated aDNA reads back to the bowhead whale reference genome using the same parameters as implemented for the ancient specimens. Finally, we downsampled the aDNA simulated individuals to 1x using SAMtools. We computed the genotype likelihoods in this simulated dataset in ANGSD using the same filtering as for the complete dataset and computed PCAs using PCAngsd. We used the sites recovered after filtering in the simulated dataset and reran a PCA with the original dataset. Finally, using the same sites uncovered in the simulated aDNA damage patterns dataset, we ran a PCA using only the subfossil individuals.

We quantified the levels of genetic divergence between pre-whaling and post-whaling bowhead whales and between localities using fixation index (*F*_ST_) values. We pooled individuals into one of four populations; pre-whaling Canada, pre-whaling Svalbard, post-whaling Canada, post-whaling Svalbard. We created a consensus pseudohaploid base call (- dohaplocall 2) file in ANGSD at the sites passing filters while computing the genotype likelihoods in the simulated dataset PCA (-sites), and using the following filters; - doMajorMinor 1 -rmtrans 1 -doMaf 2 -SNP_pval 1e-6 -minmaf 0.1 -skiptriallelic 1 - uniqueonly 1 -minind 40. We calculated *F*_ST_ in 500 kb non-overlapping sliding windows, with a minimum requirement of 100 sites per window using the available popgenWindows.py (https://github.com/simonhmartin/genomics_general).

### Genome-wide SNP heterozygosity

It has been suggested previously that heterozygosity can be estimated relatively accurately in very low-coverage individuals (<1x) when using genotype likelihoods and sites with common variants (minor allele frequency >0.1) ^131^. We tested this with our dataset by calculating heterozygosity independently, five times, on three different high-coverage ‘East Greenland Svalbard Barents Sea’ individuals, with different simulated treatments using the filtered sites obtained during the PCA analysis. Treatments included (i) no treatment (i.e. the full high-coverage dataset), (ii) the same dataset downsampled to 2x, (iii) R1 reads trimmed to 80 bp and the addition of aDNA damage patterns, (iv) R1 reads being trimmed to 80 bp, the addition of aDNA damage patterns, and downsampled to 1x, and (v) trimmed R1 reads to 80 bp, the addition of aDNA damage patterns, and downsampled to 0.2x. The simulated aDNA damage was added as described above for the PCA tests.

We calculated heterozygosity for each individual independently for the filtered sites using genotype likelihoods in ANGSD with the following parameters: -minmapQ 30 -minQ 30 -doCounts 1 -GL 2 -doMajorMinor 1 -rmtrans 1 -doMaf 2 -skiptriallelic 1 -uniqueonly 1 - doSaf 1 -fold 1 -capdepth 2. We computed a folded SFS from the sample allele frequencies using realSFS, part of the ANGSD toolsuite. To fold the SFS we used the reference genome as both the -ref and -anc parameters. To calculate the variance in our results we randomly sampled 500 thousand sites from our SNP panel 20 times, and independently calculated heterozygosity for each individual using each of the 20 subsampled SNP panels. Based on our results, we proceeded with the empirical data and restricted our heterozygosity estimates to the sites passing filters in our simulated aDNA damage PCA. We calculated the heterozygosity for all individuals in our dataset >0.2x in coverage following the same protocol. Furthermore, as our tests on the impact of aDNA damage showed a bias towards higher heterozygosity in ancient specimens, we calculated the average difference between contemporary and simulated aDNA results (0.008) and subtracted that from the values obtained for our ancient specimens.

We tested for significant differences between the pre-whaling Holocene specimens, contemporary ‘East Canada West Greenland’ stock specimens and contemporary ‘East Greenland Svalbard Barents Sea’ stock specimens by pooling the 20 subsampled heterozygosity estimates from all individuals in the given bin together and performing a Mann-Whitney-Wilcoxon Test in R v4.1.1 ^132^.

### Genome-wide nucleotide diversity

We investigated nucleotide diversity through time by splitting our dataset of individuals with >0.2x coverage into 1,000 year time bins. As the PCA suggested a single panmictic population in the pre-whaling Holocene, we kept contemporary Canada and contemporary Svalbard bowhead whales as two separate populations, but pooled the ancient individuals from the two regions to increase sample size in the pre-whaling Holocene time bins. We created a consensus pseudohaploid call (-dohaplocall 2) file in ANGSD at the sites passing filters, while computing the genotype likelihoods in the simulated dataset PCA (-sites parameter), using the following filters; -doMajorMinor 1 -rmtrans 1 -doMaf 2 -SNP_pval 1e-6 -minmaf 0.1 -skiptriallelic 1 -uniqueonly 1 -minind 40. We estimated nucleotide diversity from the pseudohaploid call file in 500 kb non-overlapping sliding windows, with a minimum requirement of 100 sites per window using the popgenWindows.py (https://github.com/simonhmartin/genomics_general). We assessed the significance of differences between the bins using a Mann-Whitney-Wilcoxon Test in R v4.1.1 ^132^. We used a Bonferroni correction to identify the threshold for significance (p-value of 0.05/6038 windows), giving us an upper p-value for significance of 0.000008.

### Simulating bottleneck impact on genetic diversity estimates

We used individual-based, forward-in-time simulations in SLiM3 ^133^ to investigate which levels of population decline and migration best fit our empirical data. We simulated two ancestral populations of different sizes based on pre-whaling estimates of the ‘East Greenland Svalbard Barents Sea’ stock (52,500 bowhead individuals) ^9^ and the ‘East Canada West Greenland’ stock (∼18,500 individuals) ^45^. We converted population size into effective population size by dividing by 10, resulting in Ne=5,250 and Ne=1,850. We simulated the populations to accumulate neutral mutations for a burn-in period of 60,000 generations until reaching mutation-drift equilibrium, in a 10 Mb genomic region with a recombination rate of 1.0×10^-08^ ^134^ and a mutation rate 2.77×10^-08^ ^135^. The ancestral populations experienced variable migration rates of either one, five or ten individuals per generation. As bowhead whales have long generation times (35-50 years, ^17^), and commercial whaling only occurred within the last ∼500 years, we investigated whether the industrial whaling bottlenecks are too recent to be visible in genetic diversity estimates of contemporary individuals. Once populations reached mutation-drift equilibrium, and assuming the final point of the simulations as the present time, we simulated bottlenecks of varying severities, 15 generations ago (or 525 years ago, assuming a conservative 35 year generation time). We calculated nucleotide diversity per population and genetic differentiation (*F*_ST_) between populations through time.

We first explored a wide range of parameters including all pairwise combinations of 20, 50, 75, 99% of population decline, variable migration rates as above, and sustained migration or interrupted migration after the bottleneck - resulting in 60 unique parameter combinations. After an initial exploratory phase, we refined the parameter space using combinations that approximately resembled the empirical data and expanded our parameter search. In a subsequent testing phase, we used a range of population decline between 40%- 90% for Canada and 90-98% in smaller step increments, assuming five migrants per generation pre-bottleneck, and either one migrant or no migration post-bottleneck, totalling 420 unique parameter combinations. For determining which simulation parameters most closely matched the empirical data, we compared the nuclear genomic nucleotide diversity of the 500-1500 bin to that of the contemporary individuals. This revealed that the contemporary ‘East Greenland Svalbard Barents Sea’ stock individuals had a nucleotide diversity proportion of 0.98 relative to the pre-whaling estimate and the contemporary ‘East Canada West Greenland’ stock individuals had nucleotide diversity proportion of 1.01 relative to the pre-whaling estimate. We also compared the heterozygosity of all fossil individuals to their contemporary counterparts resulting in proportions of 0.98 and 1.00 respectively. We also compared pre- and post-whaling *F*_ST_ values between individuals from the two different regions revealing a 3.7x increase between contemporary populations. Taking all three values into account - change in nucleotide diversity pre and post whaling for both populations, as well as change in *F*_ST_) - we selected the simulations where the population representing Canada decreased 48%, the population representing Svalbard decreased 97%, and migration between the populations ceased after the bottleneck as the top performing model and extended the simulations forward in time to estimate the predicted trajectory of genetic diversity and differentiation assuming stable post-bottleneck population sizes.

### Allele changes correlating with time

As input for Ohana ^136^, we created a genotype likelihood beagle file for all Holocene fossil individuals with >0.2x (-doGlf 2) using ANGSD and the following parameters; - minmapQ 25, -minQ 25, -uniqueonly 1. We used the GATK algorithm to call genotype likelihoods (-GL 2), calculate per-site allele frequencies assuming a fixed major and unknown minor allele (-doMaf 2), calculated major and minor alleles using GL (- doMajorMinor 1), minor allele frequency of 0.05 (-minmaf 0.05).

The genotype likelihoods of all individuals was converted to lgm as the input for Ohana using the convert function bgl2lgm. We used qpas to estimate the ancestral component proportions matrix Q (number of individuals x number of ancestral components) and allele frequencies matrix F (number of ancestral components x number of SNPs) from the genotype matrix (lgm) file with the number of ancestral components (-k) ranging from 2 to 6 with an iteration stopping criteria from log likelihood difference (-e 0.0001). In the end, we scanned for selection in each ancestral component while taking into account the sample age as a vector using neoscan.

To test for the significance of the relationship between allelic change and time, we converted the lle_ratio scores to p-values under a mixture of chi-square distributions^137^, and found the best-fitting genome-wide parameters of the mix using a Kolmogorov-Smirnov test in R. We further investigated sites with a p<0.01. Because the Holocene fossil bowhead individuals likely represented a single population, we extracted the sites with p<0.01 from each independent ancestral component run, retaining only those overlapping across all runs. We overlaid the remaining sites with the bowhead annotation using bedtools intersect ^138^ and retained any that were found within a known protein coding gene in the bowhead whale genome annotation. We BLASTed the gene sequences to find the putative gene name and used genecards.org and the NHGRI-EBI Catalog of human genome-wide association studies (https://www.ebi.ac.uk/gwas) to designate putative function. To visualise allele changes with time, we used a haploid base call (-dohaplocall 2) in ANGSD for each individual.

### Demographic history from modern genomes

We attempted to reconstruct the recent demographic history of the ‘East Greenland Svalbard Barents Sea’ stock using genetic optimization for *N*_e_ estimation (GONE)^139^ using the 12 high coverage genomes. As input we generated a PLINK file using the largest 150 autosomes in ANGSD (-doplink 2) with the following parameters -uniqueOnly 1 -GL 2 - remove_bads 1 -minMapQ 20 -minQ 20 -SNP_pval 1e-6 -skipTriallelic 1 -doMaf 2 - domajorminor 1 -minmaf 0.05 -dopost 1 -doplink 2 -minInd 12.

We ran the GONE software using two different parameter sets, one with the default parameters, but with the maximum number of SNPs per scaffold as 10000, and the other with the same parameters but with additional changes in the NGEN and NBIN parameters to 1000 as previously suggested^140^. Each of the parameter sets were run for 100 replicates. We calculated the mean and 95% confidence intervals from these replicates in R.

### Mitochondrial genomes

We generated mitochondrial genome consensus sequences for Late Pleistocene, pre-whaling Holocene and contemporary individuals using a consensus base call approach (- dofasta 2) in ANGSDv0.921^127^ for each individual independently, using the following parameters; minimum base and mapping qualities of 25 (-minmapQ 25, -minQ 25), only include reads that map to a single site uniquely (-uniqueonly 1), minimum read depth of 5 (- mininddepth 5), and build the consensus sequence only for the mitochondrial genome (-r KY026773.1). Only individuals with an average coverage of at least 10x (Late Pleistocene = 3, pre-whaling Holocene = 104, contemporary = 19) were included in further analysis. We also downloaded three mitochondrial genomes from contemporary Svalbard individuals recently confirmed to have come from unique individuals; individuals named A, H and I ^17^.

### Mitochondrial nucleotide diversity

To investigate changes in genetic diversity through the Holocene, we estimated nucleotide diversity (π) ^141^ with DnaSP v.6.12.03 ^142^ for each sampling area for every 1,000 year time bin. We pooled samples from Canada and Svalbard for the ancient specimens but kept contemporary populations separate. We excluded gaps and missing data from the analyses.

### Population structure

We constructed an unrooted haplotype network for all complete mitochondrial genomes (Late Pleistocene, pre-whaling Holocene, contemporary) individuals using the Median-joining network ^143^ as implemented in PopART ^144^. Fixation index values (*F*_ST_) were calculated by pooling individuals into 2,000 year time bins and splitting them into their two respective regions using Arlequin v3.5 ^145^, using default parameters and an input file generated with DnaSP. P-values were calculated using 1000 permutations and a significance was defined as a p-value < 0.05.

### Demographic history

We inferred the changes in female effective population size (*Ne*_(f)_) through time employing the Bayesian skyline plot method ^146^ implemented in BEAST v.2.6.1 ^147^. Based on the network (supplementary figure S9) and *F*_ST_ analyses of our 107 complete mitochondrial genomes, which spanned >30,000 years in age, we do not see any evidence for population structure, and thus we treated all the data as a single population. We aligned the mitochondrial genomes of all individuals and extracted 38 regions, including protein-coding regions, rRNAs, tRNAs and the control region, based on published coordinates. The sequences of these 38 regions were combined into six subsets, (i) first, (ii) second, and (iii) third codon position of the protein-coding regions, (iv) tRNAs, (v) rRNAs, and (vi) the control region. The best-fit partitioning scheme and substitution model for the six subsets were identified employing Partitionfinder v.2.1.1 ^148^. The best partitioning scheme and substitution models based on the corrected Akaike Information Criterion were employed as input for Beast2.

The six partitions were analysed using unlinked substitution models that had a linked genealogy and molecular clock. We used tip dates based on the mean calibrated age of each specimen. Five groups of coalescent intervals and a strict molecular clock were assumed. Posterior distributions of parameters were estimated using MCMC sampling, which consisted of 500,000 burn-in steps followed by 500 million steps, sampled at every 10,000 steps. Convergence to stationarity and mixing were assessed using Tracer v.1.7.1 ^149^ and by running an independent replicate with a different seed. Both runs converged to the same joint density or posterior. A minimum effective sample size of 400 was obtained for all the parameter estimates.

## Acknowledgements

We thank the museums - in particular the Canadian Museum of Nature CMN, Natural History Museum Oslo NHMO, and the University Museum Bergen - and the many sample collectors (incl. Anne Karin Hufthammer, Mads Peter Heide-Jørgensen, and Erik Born), whose continued efforts in collecting, curating, and preserving fossil bowhead whale specimens from the past for the future, have enabled this study. In particular, we thank Margaret Currie for her help in finding and sampling the bowhead specimens from the Canadian Museum of Nature, and Nicholas Freymueller for helping us navigate the bowhead whaling literature.

## Funding

This study was supported by the Villum Fonden Young Investigator Programme (13151) and (37352), Independent Research Fund Denmark | Natural Sciences Forskningsprojekt 1 (8021-00218B) and Sapere Aude (9064-00025B) to EDL. Contemporary bowhead sample collections from Svalbard were funded by the Norwegian Polar Institute, with grants from the Russian-Norwegian Environment Commission and the Norwegian Research Council ICE-whales programme (244488/E10). F.R. was funded by the European Research Council (ERC) under the European Union’s Horizon Europe programme (grant agreements No. 101077592 and 951385) and by a Novo Nordisk Fonden Data Science Ascending Investigator Award (NNF22OC0076816).

## Author contributions

Conceptualization, MVW, EDL; Formal analysis, MVW, SCB, HEM, AAC, JM; Investigation, MVW, AR-I, CHSO, MBS, PS; Writing – Original Draft, MVW, EDL; Writing – Review & Editing, All authors; Funding Acquisition, EDL; Resources, EDL; Supervision, FR, PS, DF, EDL.

## Competing interests

Authors have no competing interests to declare.

## Data and materials availability

Raw sequencing data will be deposited to NCBI SRA on acceptance.

## Notes

### Competing Interest Statement

The authors have declared no competing interest.

