## Supplementary information for "Four centuries of commercial whaling eroded 11,000 years of population stability in bowhead whales"

Document includes:

Supplementary results text

Supplementary figures S1-S20

Supplementary tables S1-S16

Supplementary video S1

**Supplementary results**

**Ecological niche modelling**

Ecological niche modelling (ENM) tuning and cross-validation resulted in the optimised hypervolume model having high accuracy as measured by AUC and Boyce Index. Tuning for the hypervolume parameters resulted in our final hypervolumes using a bandwidth multiplier of 1.5, four standard deviations, and a probability threshold of 0.99. 100% of our validation points were also shown to fall within the 3-dimensional boundary of our hypervolume. Our tuned ENM projections had good ability to discriminate between areas of high suitability at our withheld validation sites, relative to background samples (AUC = 0.83; Boyce index = 0.97).

Fluctuations in habitat suitability were not affected by the modelled rapid changes in fractional sea-ice cover or variation in SST, suggesting that habitat suitability was a function of all the variables in the hypervolume (Fig. 2A-C). Habitat suitability peaked in Svalbard at the onset of the Holocene, but did not peak in the Canadian Archipelago until ~10-8 kya, when average summer SST was between 1-1.5 ^o^C and summer fractional sea-ice cover was between 40% and 50%. The ENM projected a decreasing trend in average habitat suitability from 10.5 kya to ~9.5 kya in Svalbard, however, in the Canadian Archipelago habitat suitability decreased at a slower rate from 10 kya to 5kya (Fig. 2C). After these time periods, habitat suitability fluctuated slightly around a long-term mean. Analysis of the centre of gravity of habitat suitability, indicated a southward shift of up to ~2° in both subpopulations, as habitat suitability decreased (Fig. 2D). Animations of habitat suitability through time for the study region are provided as a supplementary video (Supplementary video S1).

Throughout the Holocene, suitable habitat for bowheads showed strong connectivity across much of the Arctic, including the Russian Arctic where particularly little is known about present-day bowhead whales distribution ^1^. Despite a relative lack of contemporary information, and no fossil records from these locations, bowhead whale sightings have been reported from Franz Josef Land ^2^. Further east, bowhead whales have also been observed in the western Laptev Sea ^3^, and the north of the Novosibirsk Islands Archipelago, which separates the Laptev and East Siberian seas ^4^.

Therefore, although our finding of differentiation between contemporary stocks may reflect a recent colonisation by a distinct group of bowhead whales, this is not supported by our other findings. Our habitat modelling suggests that connected suitable habitat was present across the circumpolar Arctic throughout the Holocene and up until the present day (Supplementary video S1). Coupling this with the high mobility of bowhead whales, and the fact that Canadian and Svalbard individuals represented a single population throughout the Holocene, despite sampling sites between regions being separated by several thousand km, it seems unlikely a genetically differentiated population existed to recolonise Svalbard.

**Stable isotopes**

We successfully generated bone collagen stable *δ*^13^C and *δ*^15^N isotopes from 196 Holocene specimens (Fig 1B, Supplementary table S10). We also generated *δ*^13^C and *δ*^15^N values from four Late Pleistocene individuals (1 from Canada, 3 from Svalbard), but as they are not directly comparable to the Holocene individuals, we excluded them from downstream analyses. When plotting the results through time, we observed no obvious differentiation between genetically identified females and males (Supplementary figure S4). Therefore, we did not separate the data by sex in interpretations.

**Genomic data generation**

We successfully generated palaeogenomic data from 202 Holocene fossil bowheads via shotgun sequencing (Supplementary table S11). Of these, a total of 44 individuals had coverages ranging from 0.20x-3.45x, comprising 33 from Canada and 11 from Svalbard (Supplementary table S12).

The ancient samples had typical ancient DNA damage patterns, with elevated C-T transitions on the 3-prime end of the reads, and elevated G-A transitions at the 5-prime end of the reads (Supplementary figure S16).

We further generated partial- and low-coverage genomes (0.23x - 1.24x) from seven contemporary ‘East Canada West Greenland’ stock bowhead whales, and downloaded published data from twelve contemporary ‘East Greenland Svalbard Barents Sea’ stock bowhead whales (9.96x - 20.94x) (Supplementary table S13).

**Genetic sexing**

Genetic sexing showed the Holocene dataset of 202 individuals contained 100 females, 83 males, and 19 individuals with undetermined sex. The proportions of females to males in our dataset was relatively consistent through time at ~6:4 (Supplementary figure S15), although the proportion from the Canadian samples and the Svalbard samples varied. Of the 109 specimens available from the Canadian Arctic Archipelago, we genetically identified 45 as female, 61 as male, and 3 were undetermined. Of the 93 available specimens from the Svalbard Archipelago, we genetically identified 55 as female, 22 as male, and 16 were undetermined. Information on individual genetic sexing is provided in Supplementary table S11. We did not genetically sex the contemporary samples.

For the 196 Holocene samples for which we had stable *δ*^13^C and *δ*^15^N isotope values, 94 were genetically sexed as female (43 from Canada, 51 from Svalbard), 78 were male (57 from Canada, 21 from Svalbard), and 24 were unknown (1 from Canada, 23 from Svalbard). Our four Late Pleistocene samples were identified as: Canada: 1 male, and Svalbard: 2 females, 1 undetermined. Information on the stable isotope records and associated genetic sex is provided in Supplementary table S10.

**Species identification**

All samples that yielded mitochondrial genomes were confirmed to be bowheads based on similarity to the reference genome and to each other, validating the field identifications.

**Nuclear genomes**

**Relatedness**

When considering a relatedness coefficient score (RAB) >0.125, equivalent to first cousins, as our cut off for closely related individuals, we did not find any closely related individuals in our contemporary genomes. This is in line with previous findings ^5^, that also did not find closely related individuals in the same ‘East Greenland Svalbard Barents Sea’ individuals.

**Population structure**

Principal component analysis (PCA) investigations into nuclear population structure clustered all 44 Holocene fossil samples together, regardless of geographic origin, while the 19 contemporary Canadian and Svalbard individuals formed unique clusters (Fig 3B). We retrieved similar results when downsampling the 12 contemporary Svalbard individuals to 2x, as well as simulating ancient DNA damage on the Svalbard individuals and downsampling these genomes to 1x (Supplementary figures S7-S11). When computing a PCA with only Holocene fossil samples, one individual became separated from all others on the PC1 axis. However, all principal components showed similar eigenvalues, suggesting that there was no population structure in the dataset (Supplementary figure S12). Overall, *F*_ST_ values were very low. All comparisons had mean *F*_ST_ values >0, but the distribution of *F*_ST_ values across the genomes overlapped with 0 (Supplementary Fig S6). When comparing Holocene individuals from Canada and Svalbard, we obtained a mean *F*_ST_ of 0.002. When comparing contemporary individuals from the same regions, we obtained a mean *F*_ST_ of 0.007.

**Genetic diversity**

By comparing genome-wide SNP heterozygosity levels obtained from high-quality contemporary Svalbard individuals (ID: 17-07, 17-12, and 17-19), to those from the same individual downsampled to lower coverages, and with simulated ancient DNA damage, we investigated our ability to compare the Holocene and contemporary datasets (Supplementary figure S17 and table S14). Downsampling did not appear to cause large deviations in heterozygosity. However, when simulating ancient DNA damage, we saw a noticeable increase in heterozygosity. The average deviation between contemporary and ancient simulated samples was 0.008. We did not observe any obvious associations between sequencing depth and heterozygosity estimates (Supplementary figure S18).

When binning the data into 1,000 year time bins, we observed similar levels of nuclear nucleotide diversity (pi) across the subfossil (pre-whaling) samples; for the majority of comparisons based on the pre-whaling data (samples >500 years old), we did not observe any significant differences in diversity levels, with mean values ranging from 0.362 to 0.366 (Supplementary figure S1 and table S3). However, we did observe significantly lower levels of nucleotide diversity (Supplementary table S3) in the contemporary Svalbard individuals (0.355 +/-0.035) relative to the pre-whaling time bins, and also relative to contemporary Canada (0.366 +/- 0.042). In contrast, the diversity of contemporary Canadian individuals is not significantly different from the pre-whaling time bins.

**Simulated demographic scenarios**

We first investigated the impact of broad demographic parameters on the nucleotide diversity and *F*_ST_ of two populations modelled after those from our two study regions (Supplementary Fig S3). At low levels of migration (one individual per generation) we see a difference in nucleotide diversity between regions in Holocene individuals (based on pre-whaling population size estimates) as well as relatively high *F*_ST_ between the two populations. As the migration rate increases, the population differences in nucleotide diversity disappears and *F*_ST_ values get closer to 0.

Out of our scenarios, only at high levels of population decrease (99% loss of the original population size) do we see a decrease in nucleotide diversity in the modelled populations. Moreover, even when migration continues at comparable levels after the bottleneck (five or ten individuals per generation) we see an increase in *F*_ST_. This suggests that a decrease in population size alone is sufficient to cause increased *F*_ST_ values. However, when there is migration, nucleotide diversity decreases relatively less than when migration was completely stopped.

When conducting more fine-scale simulations, several population size decreases produce *F*_ST_ increases and nucleotide diversity decreases similar to in the empirical data (Fig 4).

**Genes under selection**

After filtering for sites with p-values < 0.01 across all K values, we were left with 140 sites. 18 sites were located within 16 annotated genes. After manual inspections for functions using the GWAS databases, we noticed the genes have been associated with a wide array of phenotypes, some of which could putatively be linked to Arctic adaptations (Supplementary table S6). Allele frequencies may change after strong positive selection, resulting in the near replacement of one allele with the selected variant. Alternatively, relaxation of previously strong selective pressure on an allele can lead to the prevalence of both alleles in the gene pool. In most cases, we see that an alternative allele becomes fixed after ~3 kya suggesting positive selection. However, in the contemporary ‘East Greenland Svalbard Barents Sea’ stock, regardless of site, the original allele is still segregating, at varying ratios, indicating that our results might reflect a mixture of both scenarios. None of these genes overlapped with a previous study looking at genes under positive selection in the bowhead whale lineage as a whole, which mostly included a number of ageing and cancer-associated genes ^6^.

All sites within the genes investigated are biallelic in contemporary individuals from the ‘East Greenland Svalbard Barents Sea’ stock (Supplementary table S15), suggesting standing variation in these genes is still present for selection to act upon. In contemporary individuals from the ‘East Canada West Greenland’ stock, five sites contain only one allele, which could limit future adaptability of these genes (Supplementary table S15). However, this finding may reflect smaller sample size (n = 7 *vs* n = 12) and lower genomic sequencing coverage for this population (Supplementary table S12). Although the genes uncovered by our selection analysis appear to have been important to buffer the impacts of past environmental change, the sites identified as showing signals of selection may not be the same sites that will be selected upon in the future. Hence retaining higher levels of genome-wide diversity, and therefore having many substrates for selection to act upon, will likely be key in enabling future adaptation.

**Mitochondrial genomes**

**Data generation**

We generated a total of 107 complete mitochondrial genomes (>10x) from the fossil material. All 65 Canadian mitogenomes and 20 of the Svalbard mitogenomes were obtained via shotgun sequencing. An additional 22 Svalbard mitogenomes were obtained via target enrichment. The mean coverages of the mitogenomes ranged between 10x and 271.45x (Supplementary tables S11, S12, S16).

We were also able to generate mitochondrial genomes from three of our four Late Pleistocene samples: CGG_1_023629 (Supplementary table S11), CGG_1_023776 (Supplementary table S12), CGG_1_023745 (Supplementary table S16).

Our seven contemporary Canadian samples all yielded complete mitochondrial genomes (496.8-3902.5x). Combining this with 15 available Svalbard samples^5,7^, we had a total sample size of 110 pre-whaling and 22 contemporary mitogenomes.

**Population structure**

Haplotype network analyses did not show any spatial or temporal subdivision (Supplementary figure S19). Seven haplotypes were shared either across space or across time; four haplotypes were shared between Canada and Svalbard, and one haplotype was shared among three Canadian individuals spanning 12,000 years. The three Late Pleistocene individuals fit within the diversity of Holocene/contemporary individuals. Investigations into Fst values compared between regions and by pooling individuals into 2,000 year time bins revealed no significant differences.

**Genetic diversity**

All time bins except contemporary Svalbard (haplotype diversity of 0.993) had haplotype diversities equal to 1 (Supplementary table S4). Furthermore, unlike with the nucleotide diversity estimates based on the nuclear data, we observed fluctuations in levels of mitochondrial nucleotide diversity through time in the mitochondrial data, albeit with highly overlapping standard deviations (Supplementary figure S1B). We observed the lowest levels of nucleotide diversity between 5-4 kya and 2-1 kya. Contemporary individuals do not show lower levels of mitochondrial nucleotide diversity than the Holocene individuals.

**Demographic history**

Our skyline plot calculated using the entire mitochondrial genome dataset of 129 individuals including Late Pleistocene, pre-whaling Holocene, and contemporary individuals show maternal effective population size (N_ef_) across the past 50 ky (Supplementary figure S20). N_ef_ was relatively low and stable until the onset of the Last Glacial Maximum ~24 kya, when we observe a rapid and continual increase until ~14 kya. After the peak in N_ef_ ~14 kya, there is a gradual decline until the present, with estimated N_ef_ 3-fold higher than estimates for the last glacial period.

**Supplementary figures**

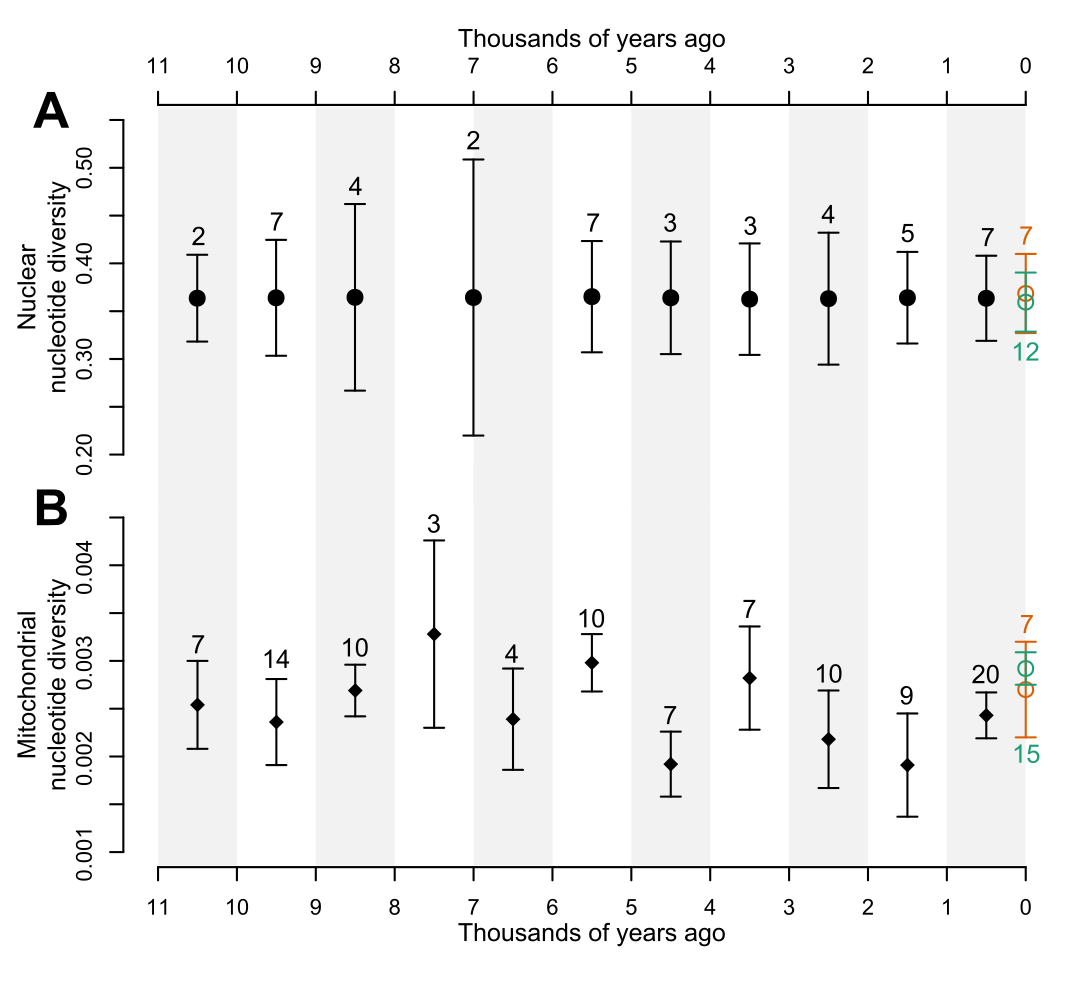

**Supplementary figure S1.** Holocene nucleotide diversity in bowhead whales. Mean (A) nuclear and (B) mitochondrial diversity are provided in 1,000 year time bins, to the exclusion of the nuclear estimates of 6,000-8,000 years ago, which were pooled into a 2,000 year time bin due to low sample size. Error bars show one standard deviation on each side of the mean. Black dots show pooled Canadian and Svalbard individuals, contemporary ‘East Canada West Greenland’ stock is in orange and the contemporary ‘East Greenland Svalbard Barents Sea’ stock is in green. Sample sizes for each bin provided. Individuals with radiocarbon dates that could not be accurately calibrated due to the marine-reservoir effect were added to the 0-1,000 bin. An overview of the estimates is also provided in Supplementary tables S2 and S4.

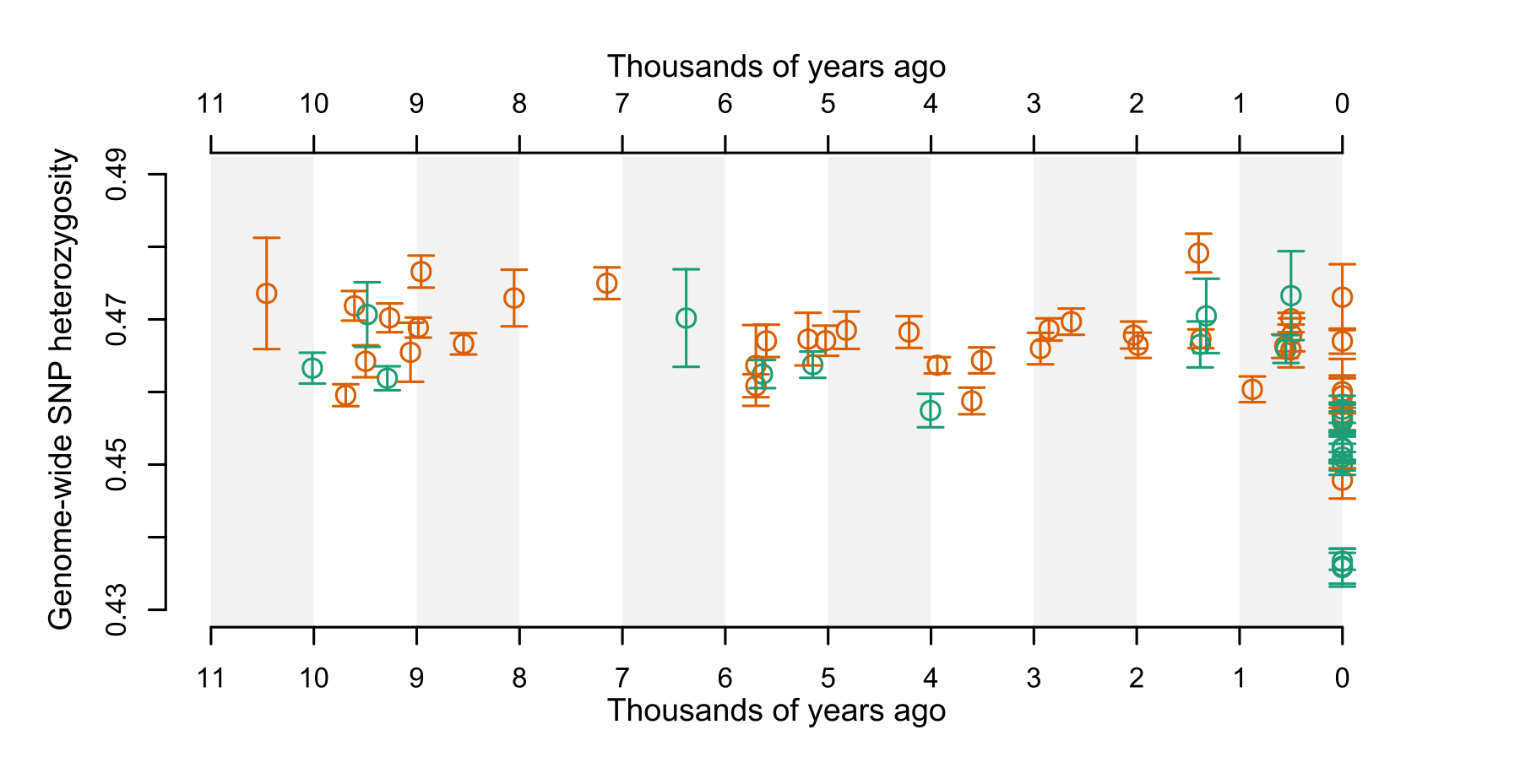
**Supplementary figure S2.** ‘Uncorrected’ individual genome-wide nuclear SNP heterozygosity. Samples from the Canadian Arctic Archipelago (n = 33 pre-whaling, 7 contemporary) are shown in orange, and from the Svalbard Archipelago (n = 11 pre-whaling, 12 contemporary) in green.

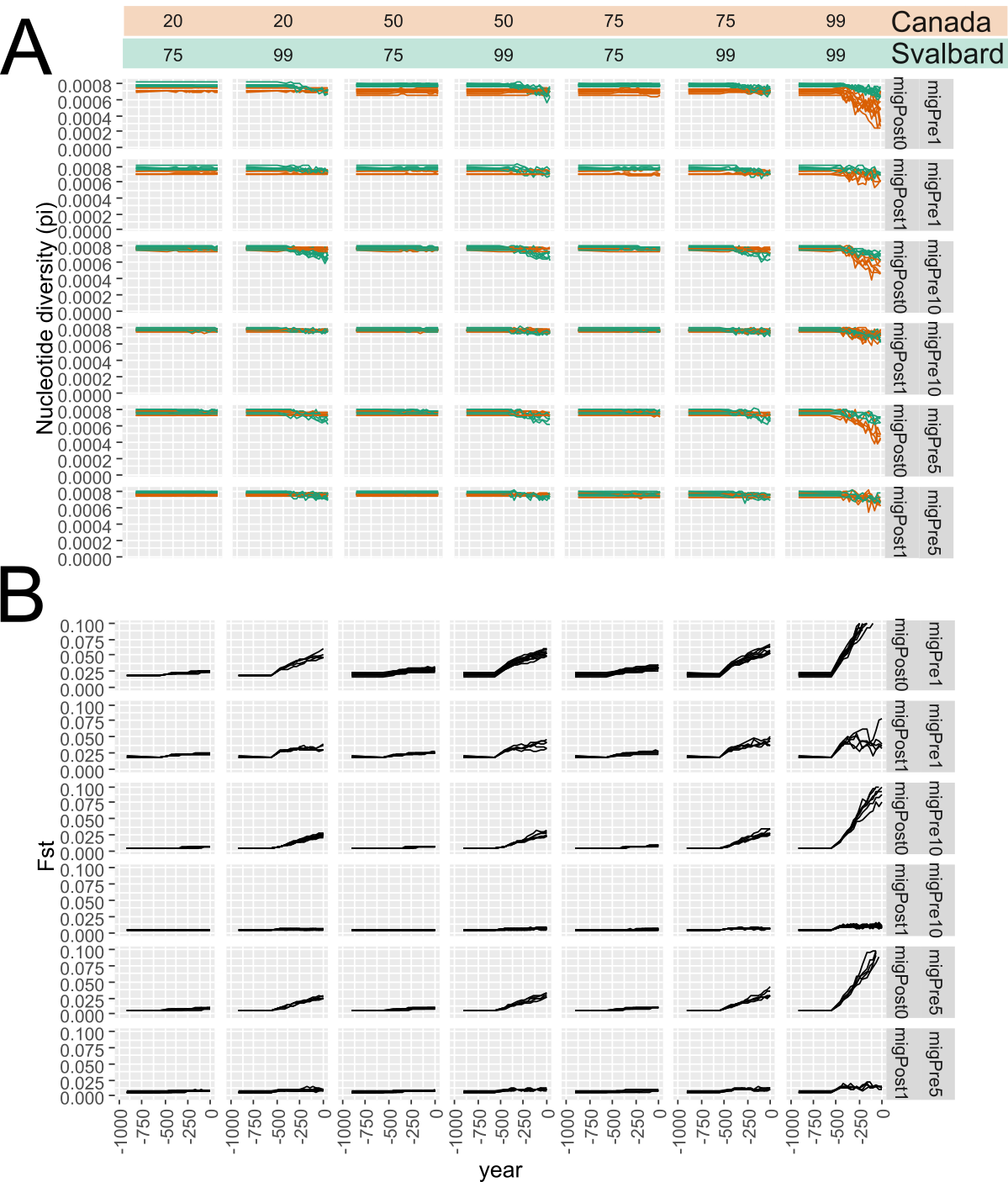

**Supplementary figure S3.** SLiM simulations exploring a broad parameter space. (A) Nucleotide diversity (pi); (B) *F*_ST_. Numbers at the top represent the percentage of population size decrease in each respective region 500 years ago (-500). migPre - number of migrants per generation prior 500 years ago, migPost - whether migration continues (1) or stopped (0) after 500 years ago.

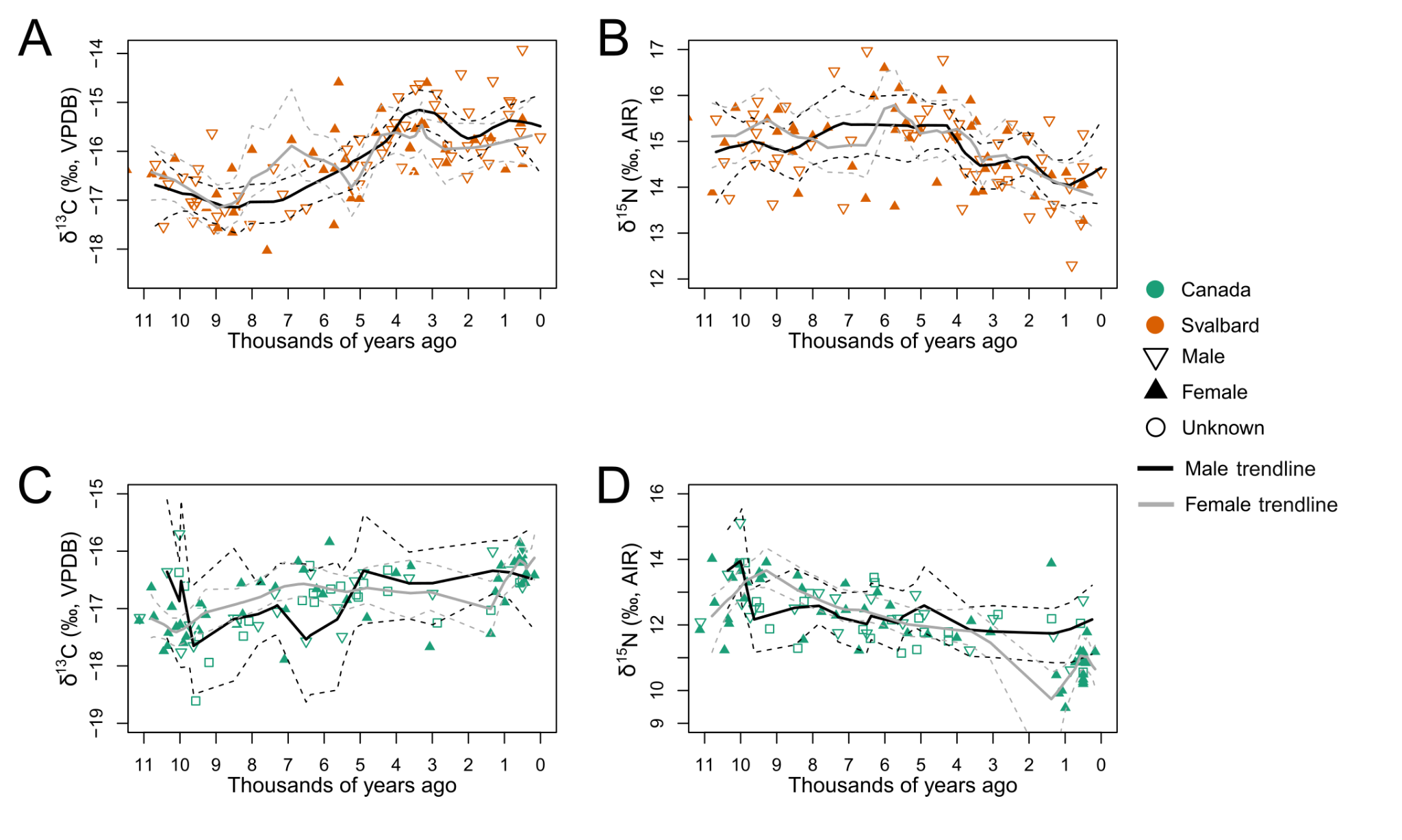

**Supplementary figure S4.** Spatiotemporal patterns of bone collagen stable *δ*^13^C and *δ*^15^N isotope values from 196 Holocene bowhead whale fossils, by sex and region. (A) *δ*^13^C values of Canadian Holocene individuals, (B) *δ*^15^N values of Canadian Holocene individuals, (C) *δ*^13^C values of Svalbard Holocene individuals, (D) *δ*^15^N values of Svalbard Holocene individuals. Sample sizes are: Canadian male n = 57, female n=43, unknown n=1; Svalbard male n = 21, female n=53, unknown n=24. Trend lines are from a local weighted regression smoothed to fit our scatterplot data and separated by sex. Stable isotope and associated genetic sexing information is provided in Supplementary table S10.

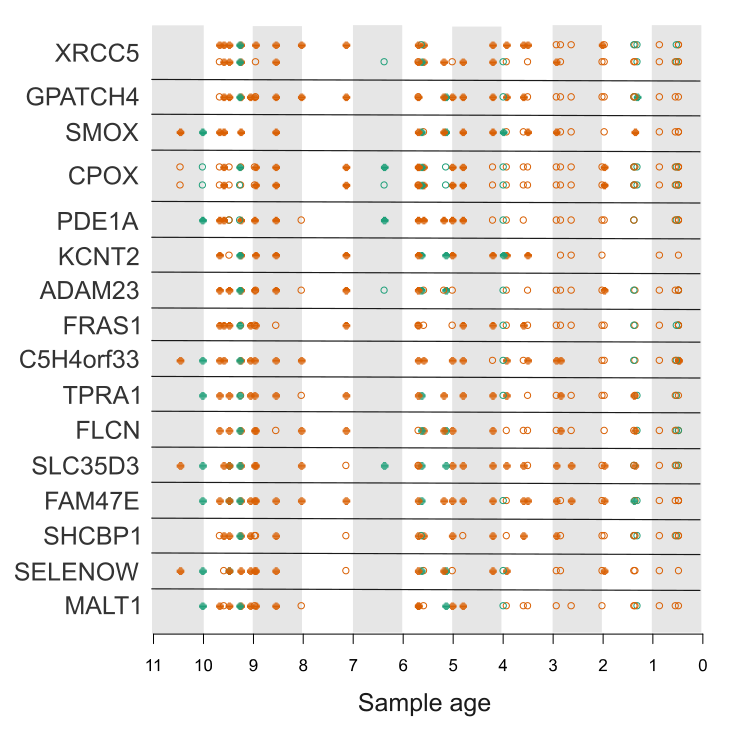

**Supplementary figure S5.** Patterns of allelic change in 18 sites across 16 genes found to have allele frequency changes significantly correlated with time, when analysing the genomes of 44 pre-whaling, Holocene fossil individuals with a genome-wide coverage >0.2x. Each circle represents the pseudohaploid state of a single individual. Open circles indicate that an individual has one copy of the two possible alleles (allele one), while the closed circles indicate the individual has the other allele (allele two). Samples from the Canadian Arctic Archipelago are shown in orange, and from the Svalbard Archipelago in green.

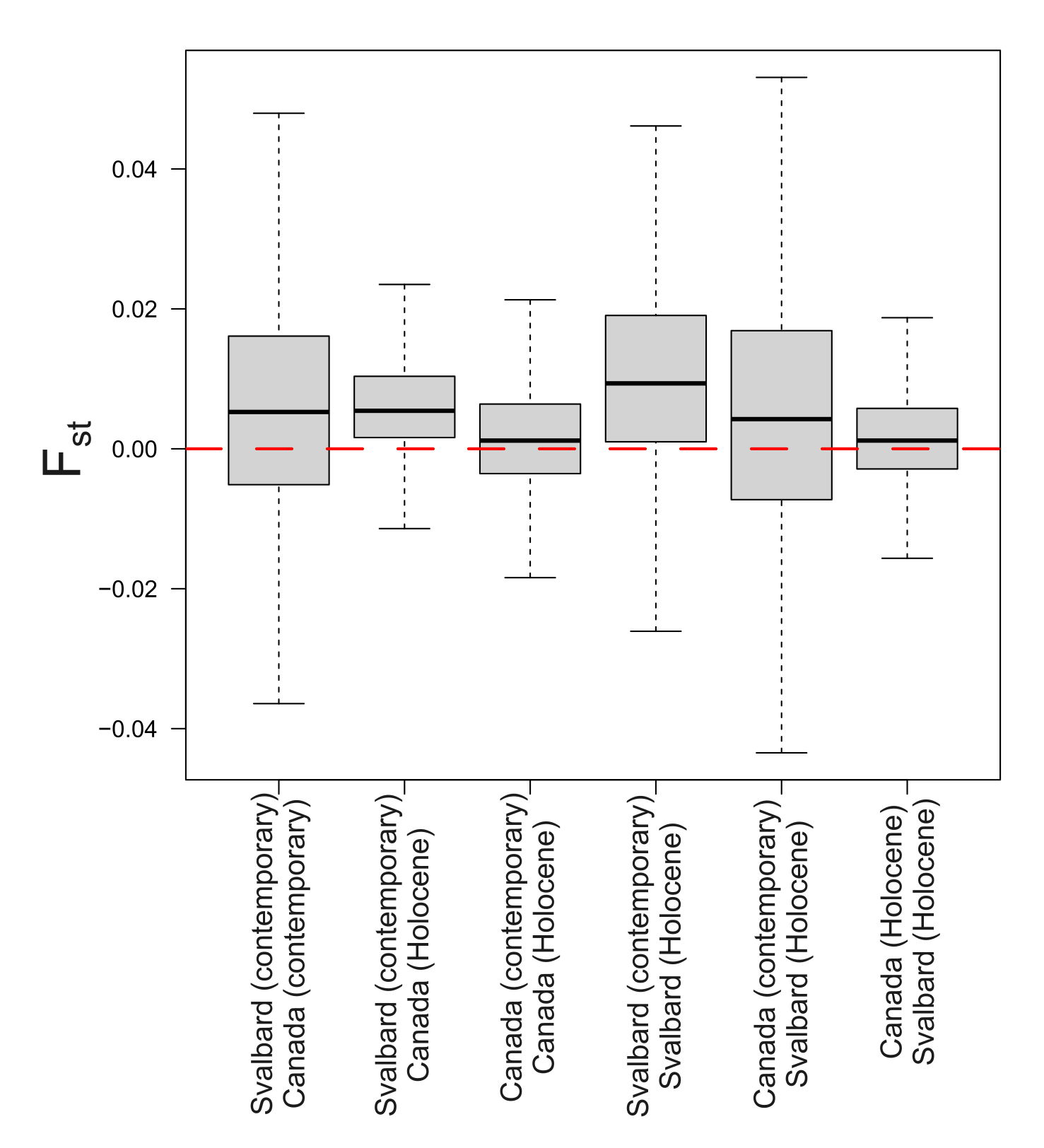

**Supplementary figure S6.** Nuclear genome-wide *F*_ST_, calculated by pooling individuals into one of four populations; pre-whaling, Holocene Canada (n = 33), pre-whaling, Holocene Svalbard (n = 11), post-whaling, contemporary Canada (n = 7), post-whaling, contemporary Svalbard (n = 12). Variation obtained by calculating *F*_ST_ in 500 kb non-overlapping sliding windows.

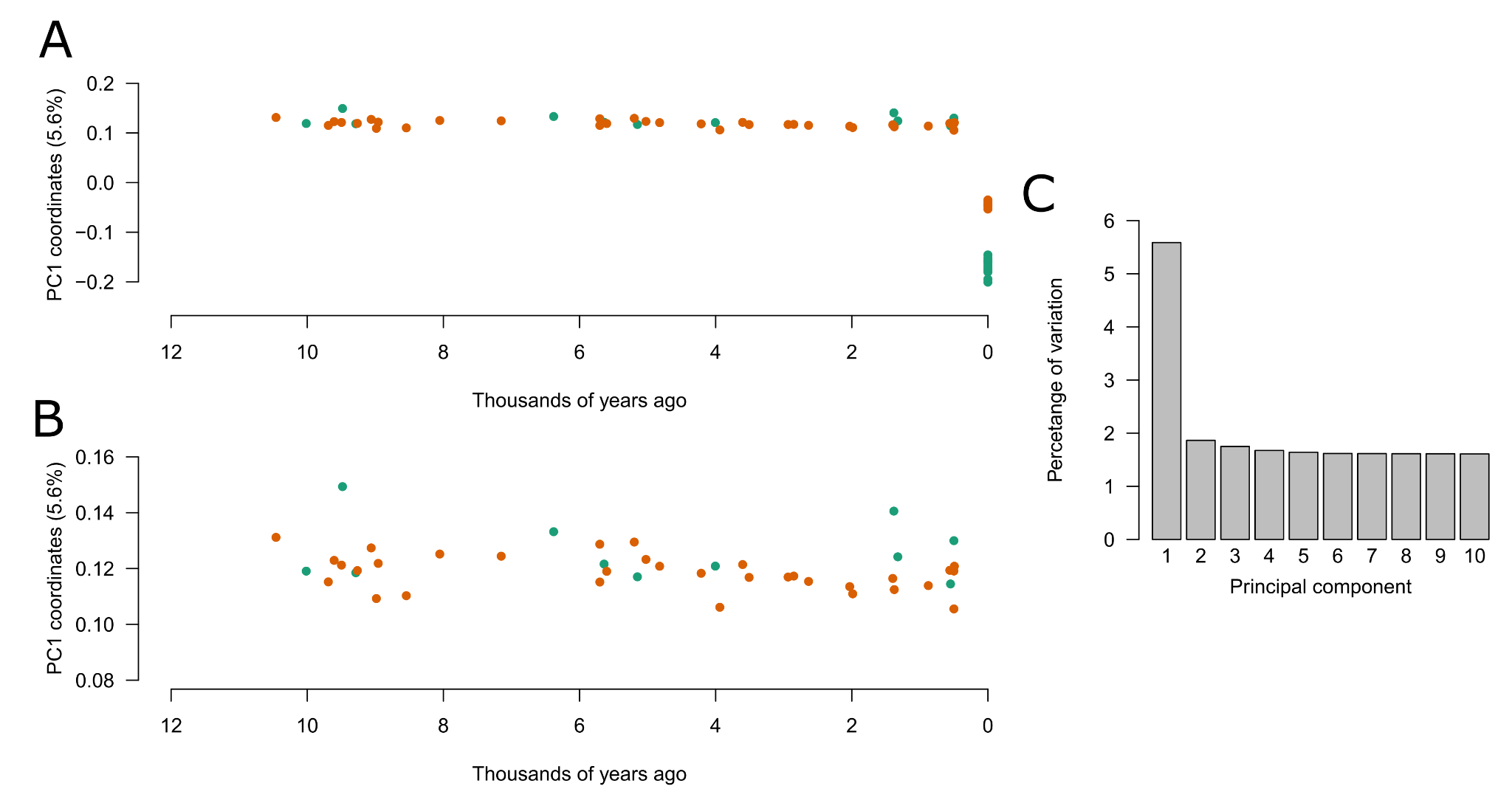

**Supplementary figure S7.** Principal component analysis using the full dataset (i.e. no downsampling or simulated ancient DNA damage) of all 44 pre-whaling fossils (>0.2x) and 19 contemporary bowhead individuals. (A) PC1 vs time. (B) PC1 vs time replotted to focus on the pre-whaling fossil individuals. (C) Percentage of the variation explained by principal components 1 to 10.

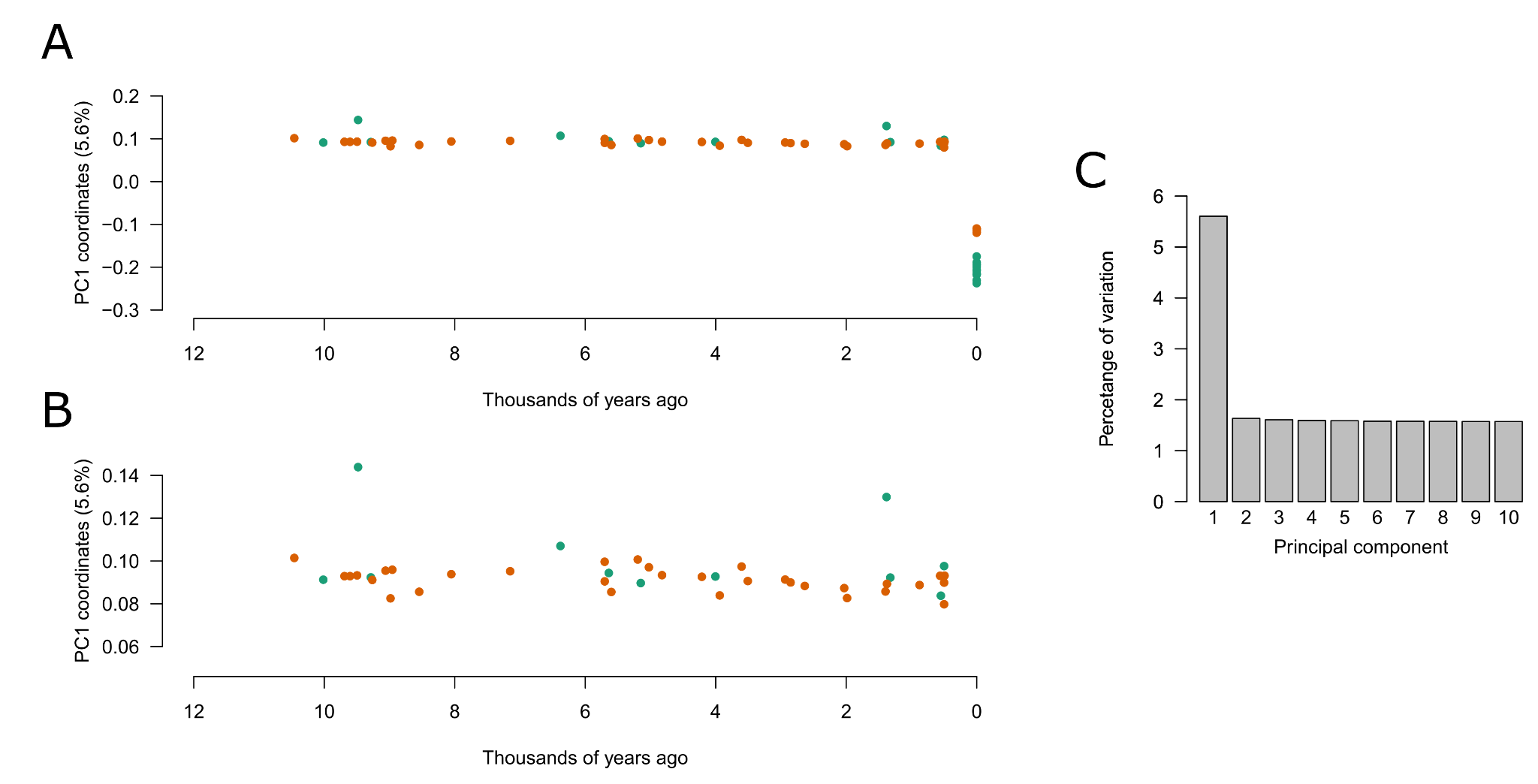

**Supplementary figure S8.** Principal component analysis using all 44 pre-whaling fossils (>0.2x) and 19 contemporary bowhead individuals. However, the contemporary ‘East Greenland Svalbard Barents Sea’ individuals were downsampled to ~2x coverage. (A) PC1 vs time. (B) PC1 vs time replotted to focus on the pre-whaling fossil individuals. (C) Percentage of the variation explained by principal components 1 to 10.

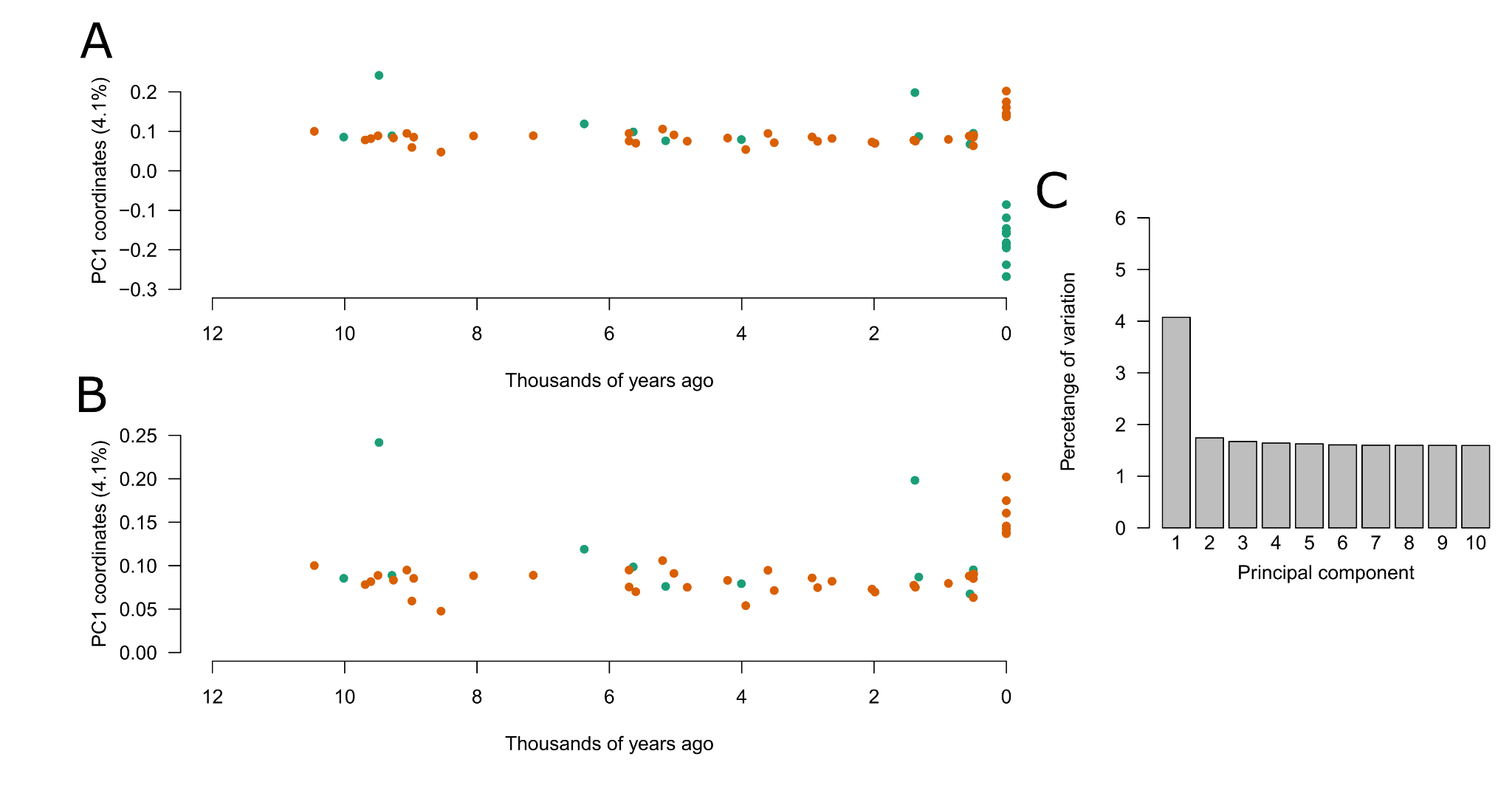

**Supplementary figure S9.** Principal component analysis using all 44 pre-whaling fossils (>0.2x) and 19 contemporary bowhead individuals. However, the contemporary ‘East Greenland Svalbard Barents Sea’ individuals have ancient DNA damage patterns simulated onto the reads as well as and reduced read lengths (80bp). (A) PC1 vs time. (B) PC1 vs time replotted to focus on the pre-whaling fossil individuals. (C) Percentage of the variation explained by principal components 1 to 10.

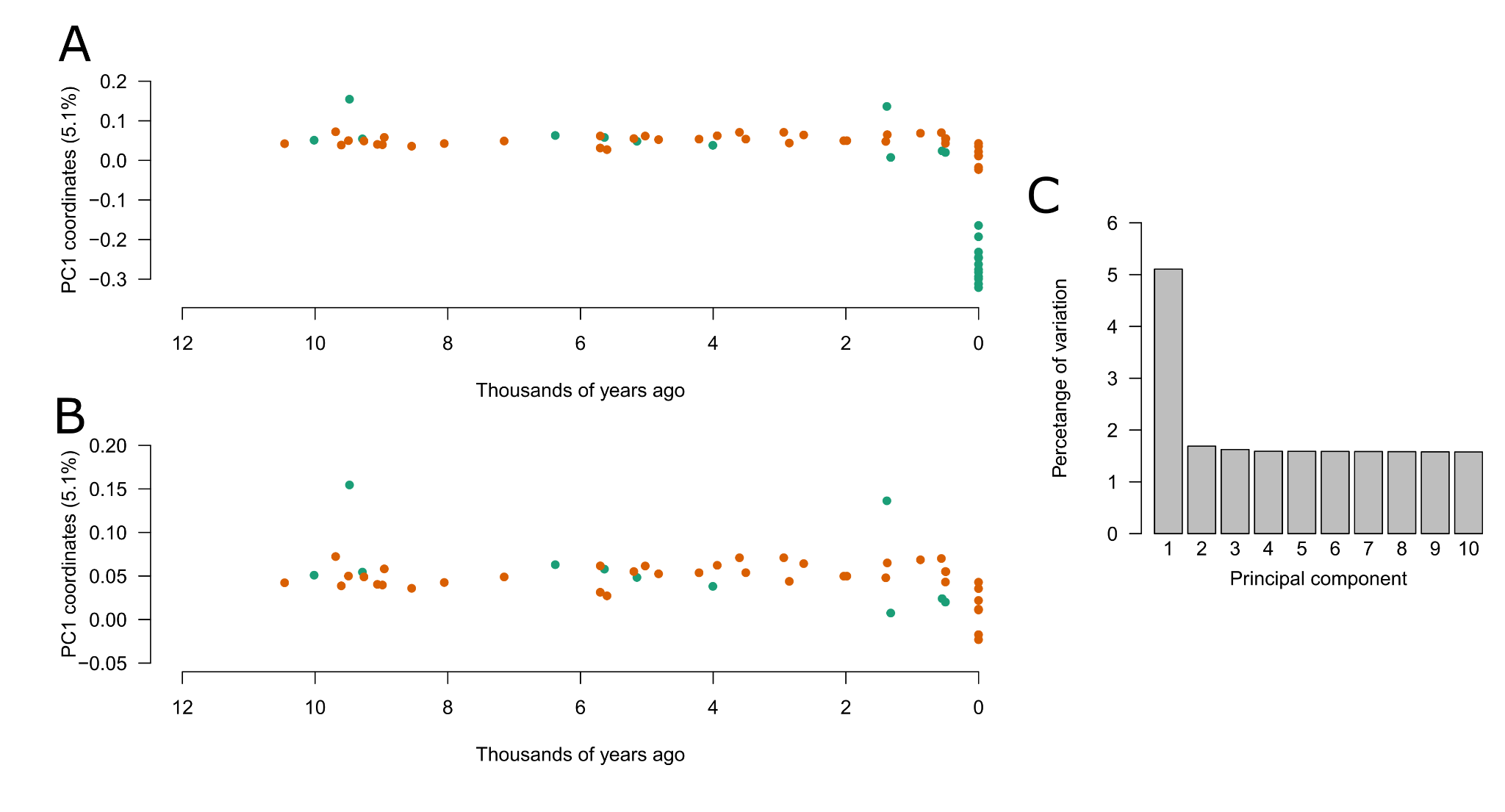

**Supplementary figure S10.** Principal component analysis using all 44 pre-whaling fossils (>0.2x) and 19 contemporary bowhead individuals. However, the contemporary ‘East Greenland Svalbard Barents Sea’ individuals have ancient DNA damage patterns simulated onto the reads, reduced read lengths (80bp), and were downsampled to 1x. (A) PC1 vs time. (B) PC1 vs time replotted to focus on the pre-whaling fossil individuals. (C) Percentage of the variation explained by principal components 1 to 10.

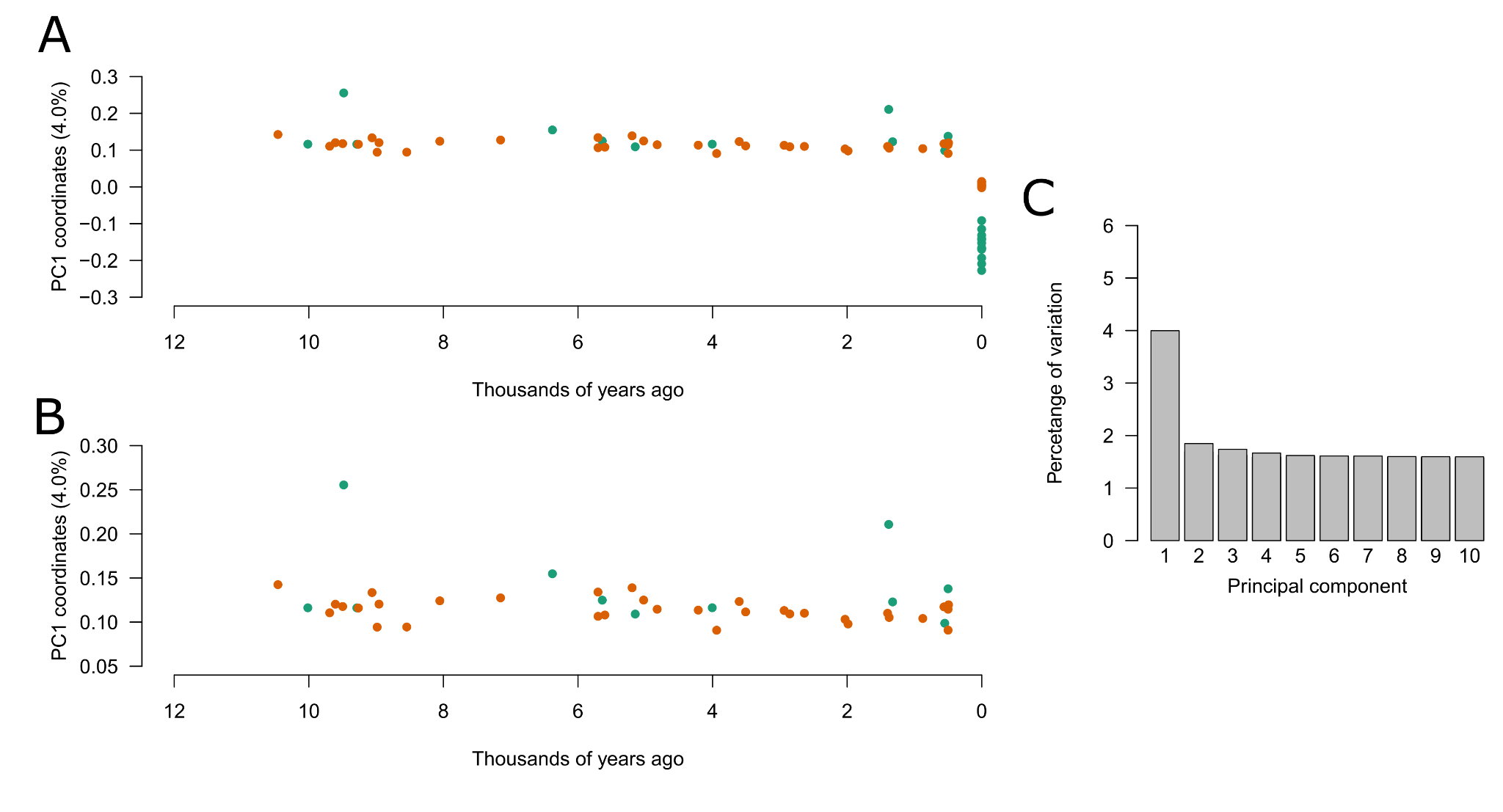

**Supplementary figure S11.** Principal component analysis using the full dataset (i.e. no downsampling or simulated ancient DNA damage) of all 44 pre-whaling fossils (>0.2x) and 19 contemporary bowhead individuals. However, this analysis only included the sites that were obtained after filtering using the data set with the contemporary ‘East Greenland Svalbard Barents Sea’ individuals with ancient DNA damage patterns simulated onto the reads as well as and reduced read lengths (80bp). (A) PC1 vs time. (B) PC1 vs time replotted to focus on the pre-whaling fossil individuals. (C) Percentage of the variation in the PCA explained by principal components 1 to 10.

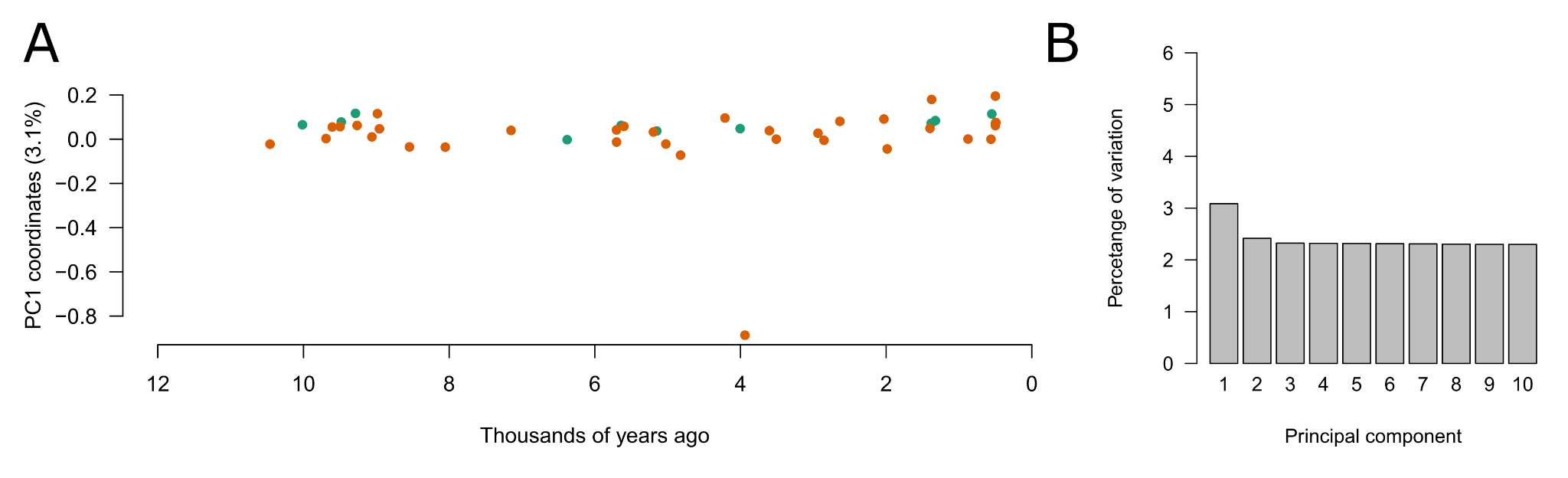

**Supplementary figure S12.** Principal component analysis using only the 44 pre-whaling fossils (>0.2x). (A) PC1 vs time. (B) Percentage of the variation in the PCA explained by principal components 1 to 10.

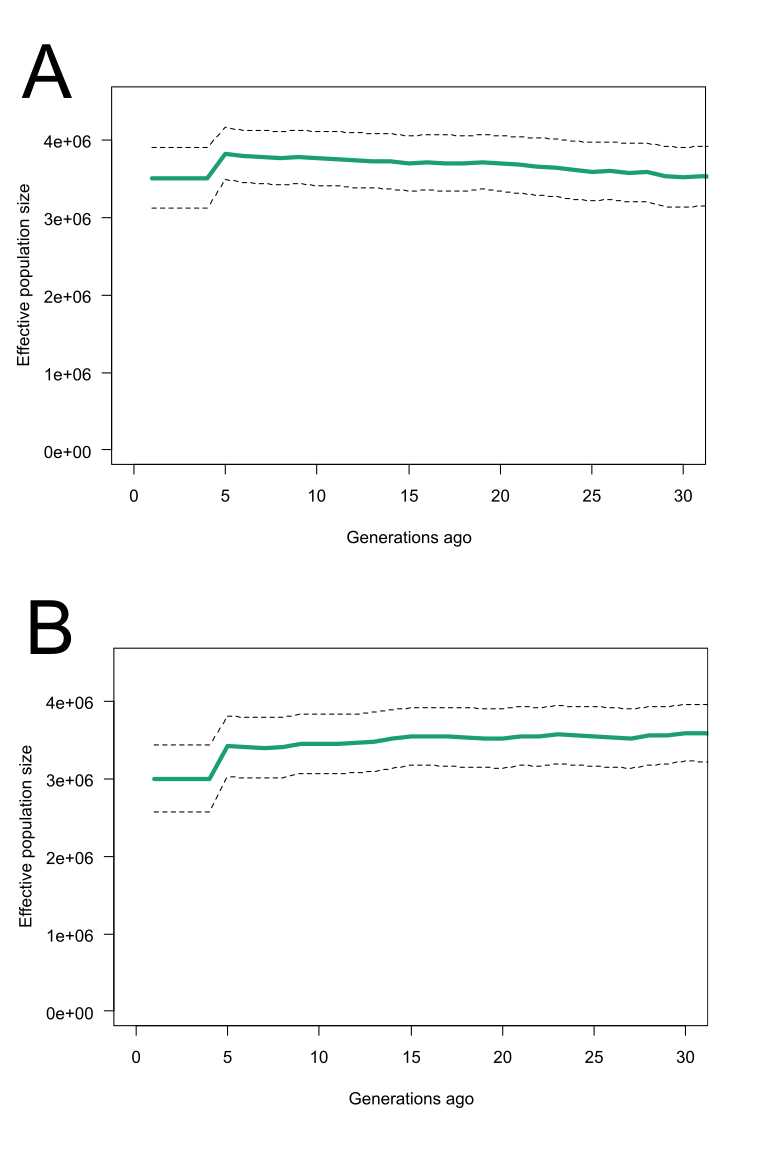

**Supplementary figure S13.** Demographic history of the ‘East Greenland Svalbard Barents Sea’ stock reconstructed with GONE, based exclusively on contemporary individuals. (A) Using default parameters, and (B) using the parameters from Kardos et al^8^. Dotted line shows the 95% confidence intervals.

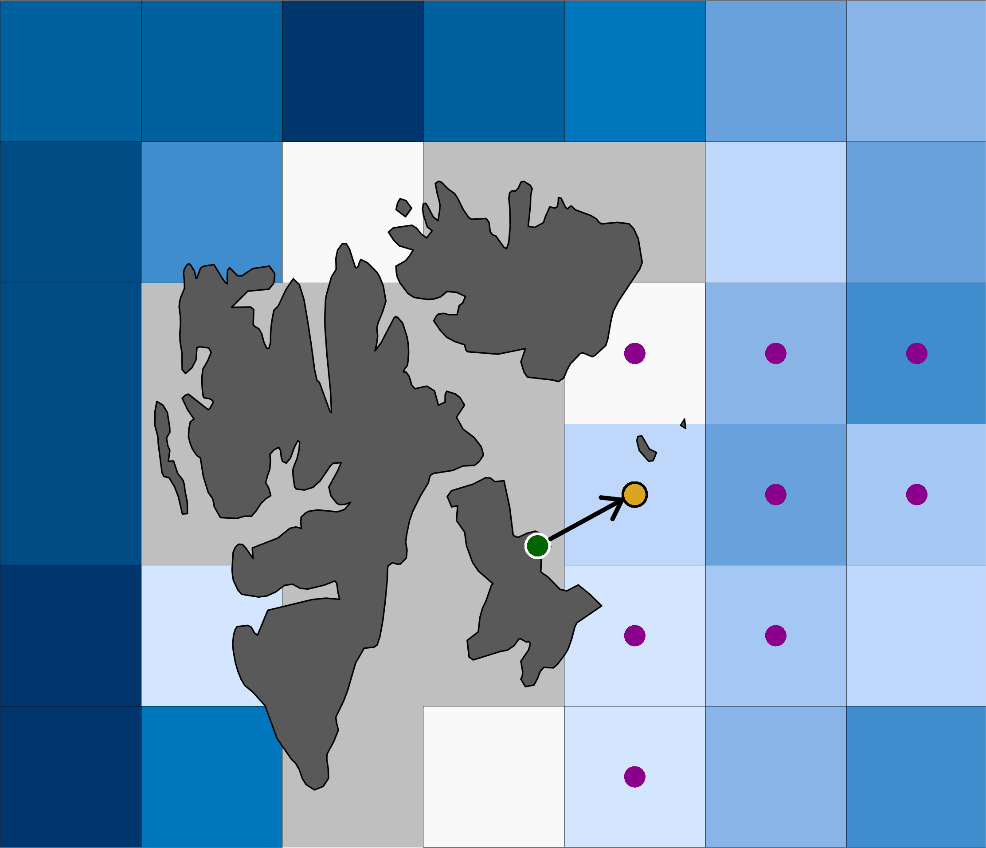

**Supplementary figure S14:** The 9-cell averaging approach used to characterise climatic and environmental conditions for each fossil site. The original fossil location (green point) is on a land-cell (grey grid-cells), this location is snapped to the nearest ocean-cell (blue grid-cells). The new location (yellow point) is then used to identify the eight closest cells (purple points). Climatic and environmental conditions are then characterised as the average conditions over these 9-cells.

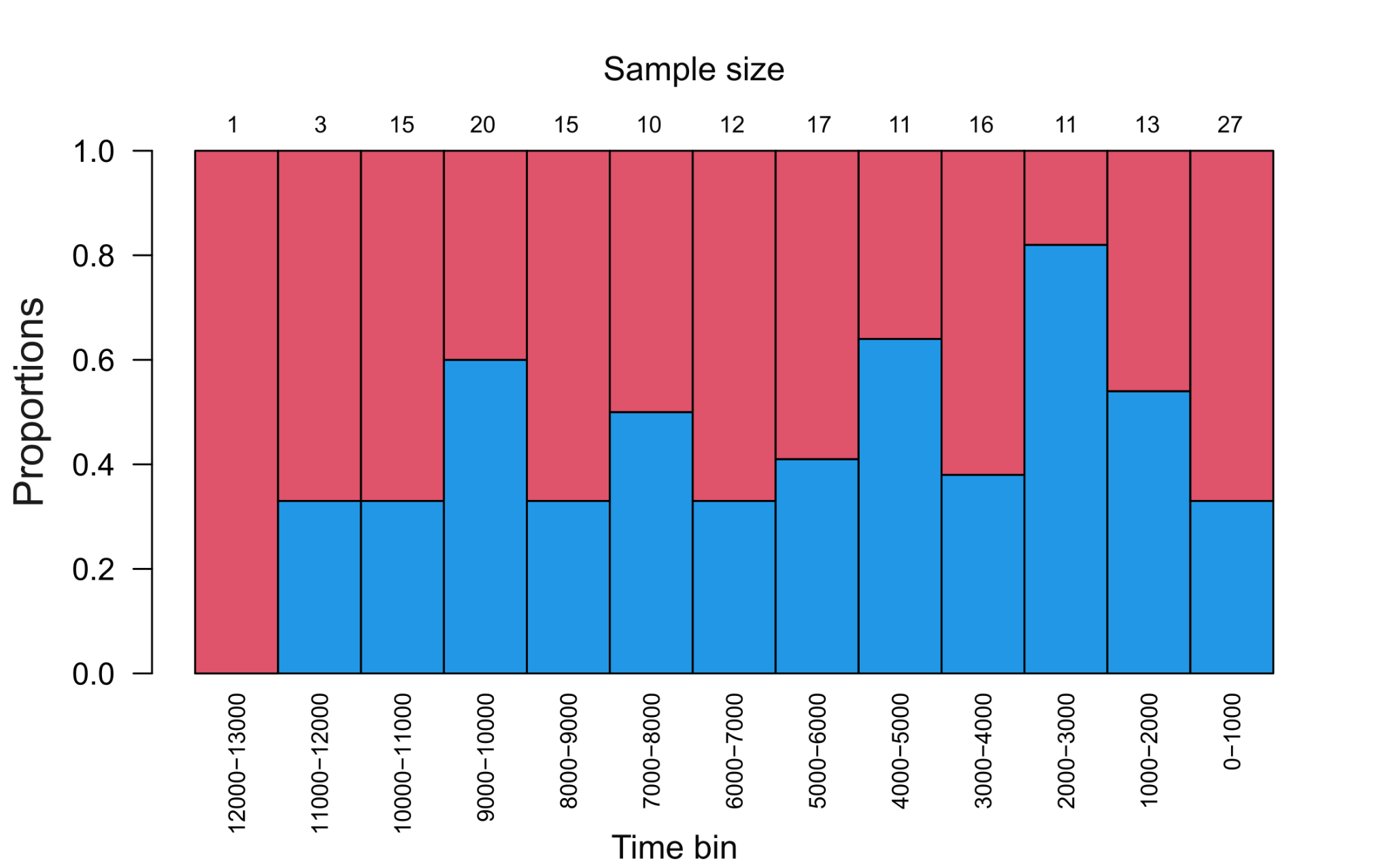

**Supplementary figure S15.** Genetic sexing of the pre-whaling, Holocene fossils. Proportions of genetically identified females (red, total n = 100; 45 from Canada, 55 from Svalbard) and males (blue, total n = 83; 61 from Canada, 22 from Svalbard) in 1,000 year time bins, across regions. We were unable to genetically identify the sex of 19 specimens (3 from Canada, 16 from Svalbard). Sample size for each bin is indicated at the top of each bar. Contemporary individuals are not shown.

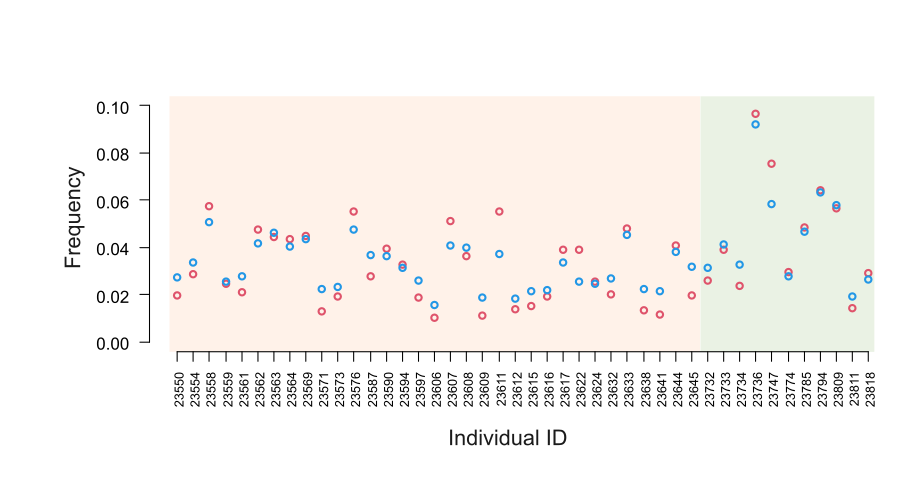

**Supplementary figure S16:** DNA damage results taken from Mapdamage for the 44 pre-whaling fossils with >0.2x coverage. The frequency of C-T transitions on the first site from the 3-prime end of the read is shown in red, and G-A transitions on the first site from the 5-prime end of the read is shown in blue. Samples from the Canadian Arctic Archipelago are in orange (n = 33), and from the Svalbard Archipelago in green ( n = 11).

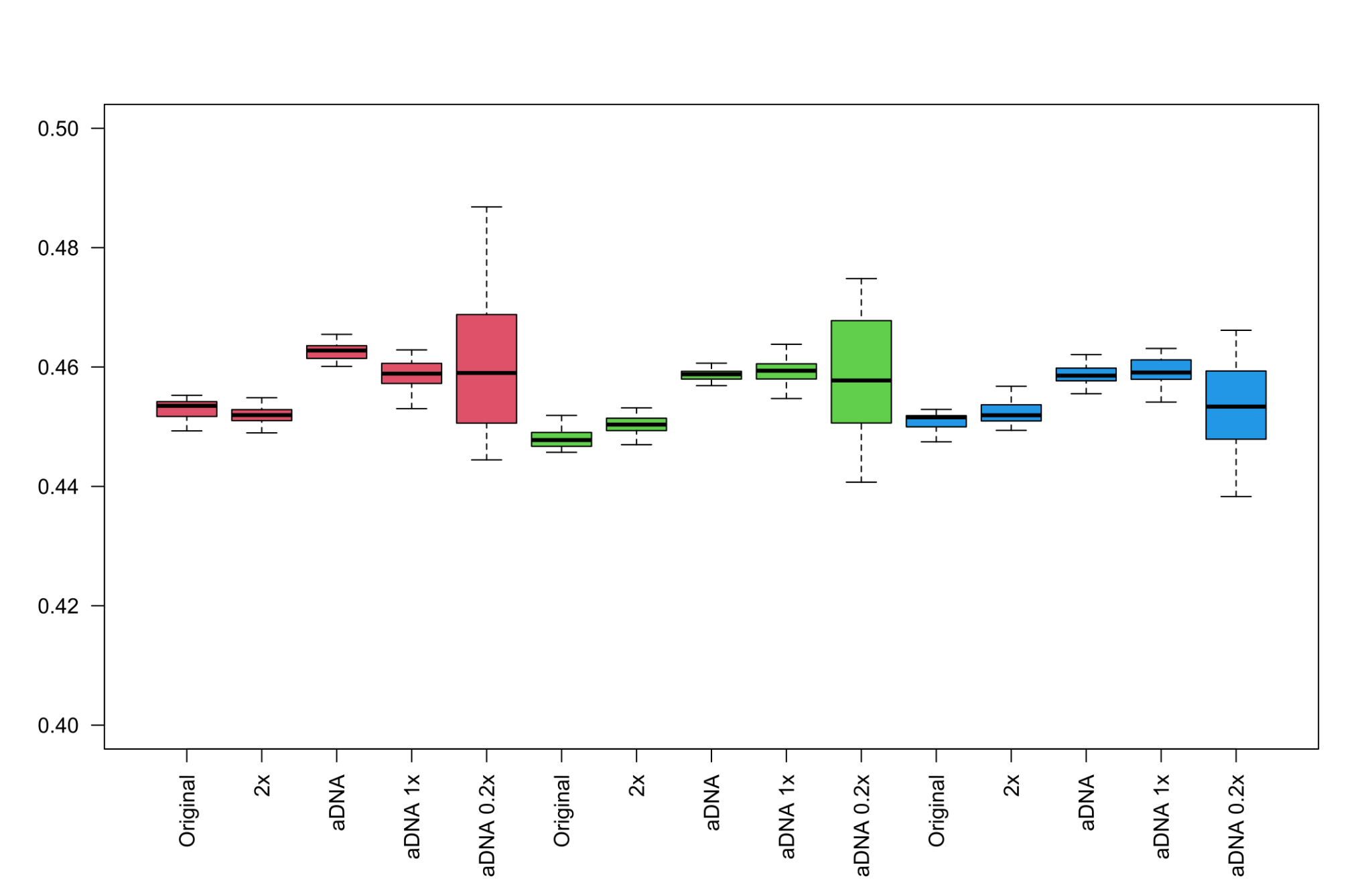

**Supplementary figure S17.** SNP heterozygosity estimates of three high-coverage individuals from Svalbard, with downsampling and simulated ancient DNA damage patterns. Heterozygosity was estimated from 500k sites randomly selected from the SNP panel 20 times independently. Individuals are colour coded. Colours correspond to individual ID; red = 17-07, green = 17-12, blue = 17-19.

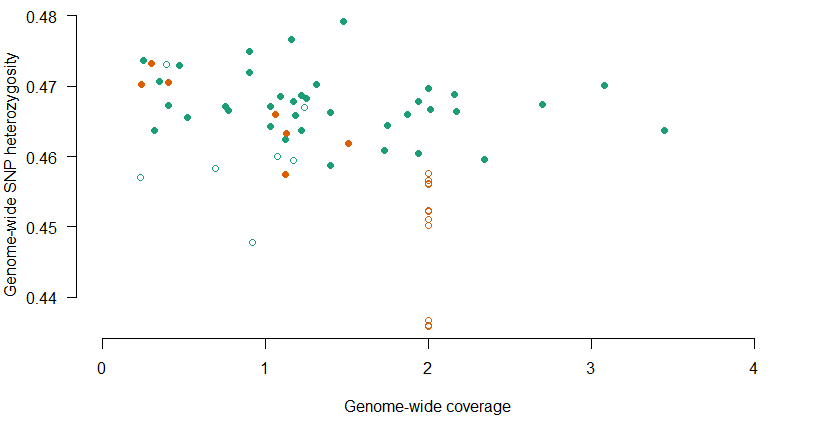

**Supplementary figure S18.** Genome-wide SNP heterozygosity estimates versus genome-wide coverage across pre-whaling, Holocene fossils (dots, n = 44) and contemporary (open circles, n = 19) bowhead whale individuals. Green indicates samples from the Canadian central Arctic and orange indicates samples from Svalbard. Only individuals with >0.2x were included in the comparison.

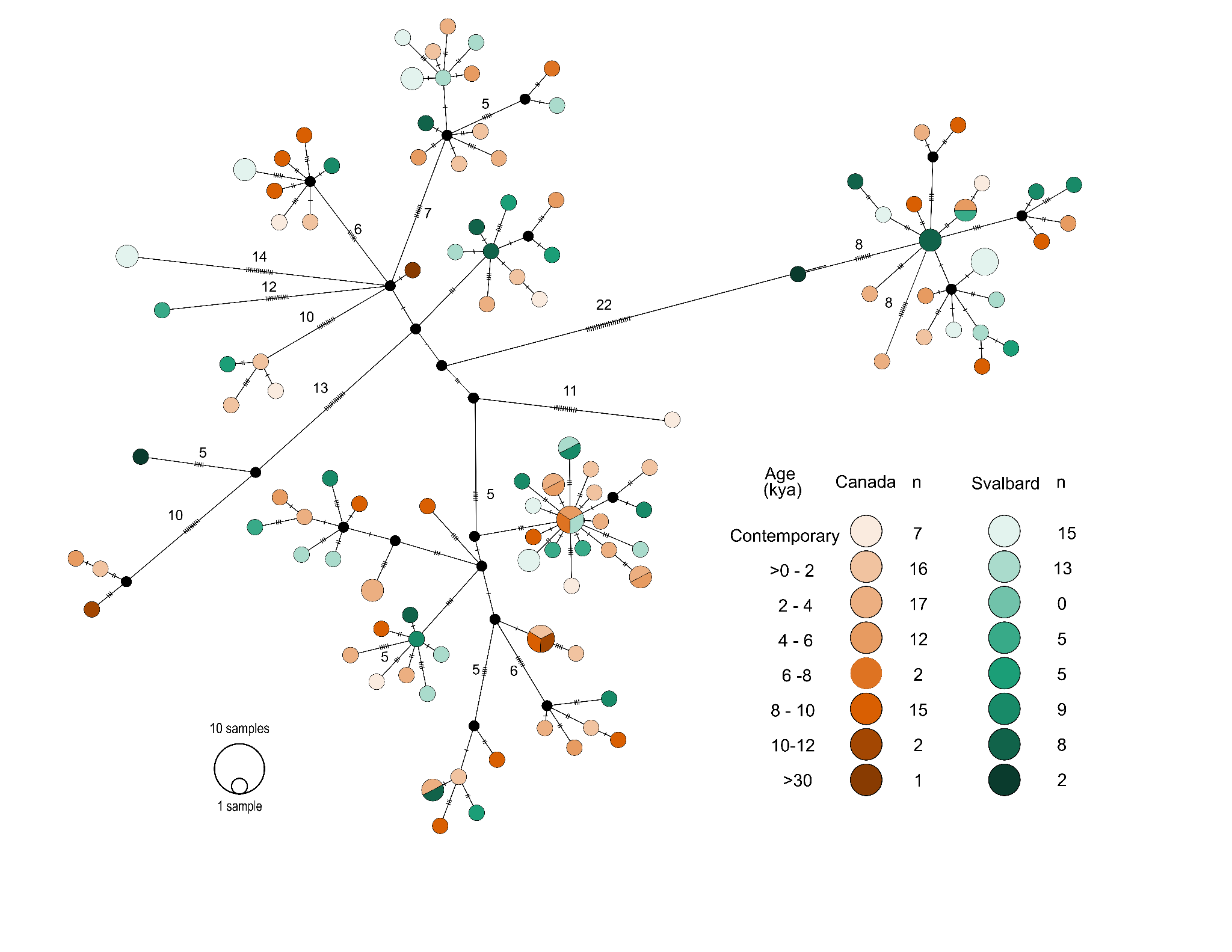

**Supplementary figure S19.** Haplotype network of 129 bowhead whale mitochondrial genomes (>10x), coloured by age bin. Sample sizes are provided, and included 65 pre-whaling, Holocene fossil sequences from Canada and 42 from Svalbard, in addition to Late Pleistocene (>30 kya) and contemporary sequences. Samples from the Canadian Arctic Archipelago are shown in orange, and from the Svalbard Archipelago in green.

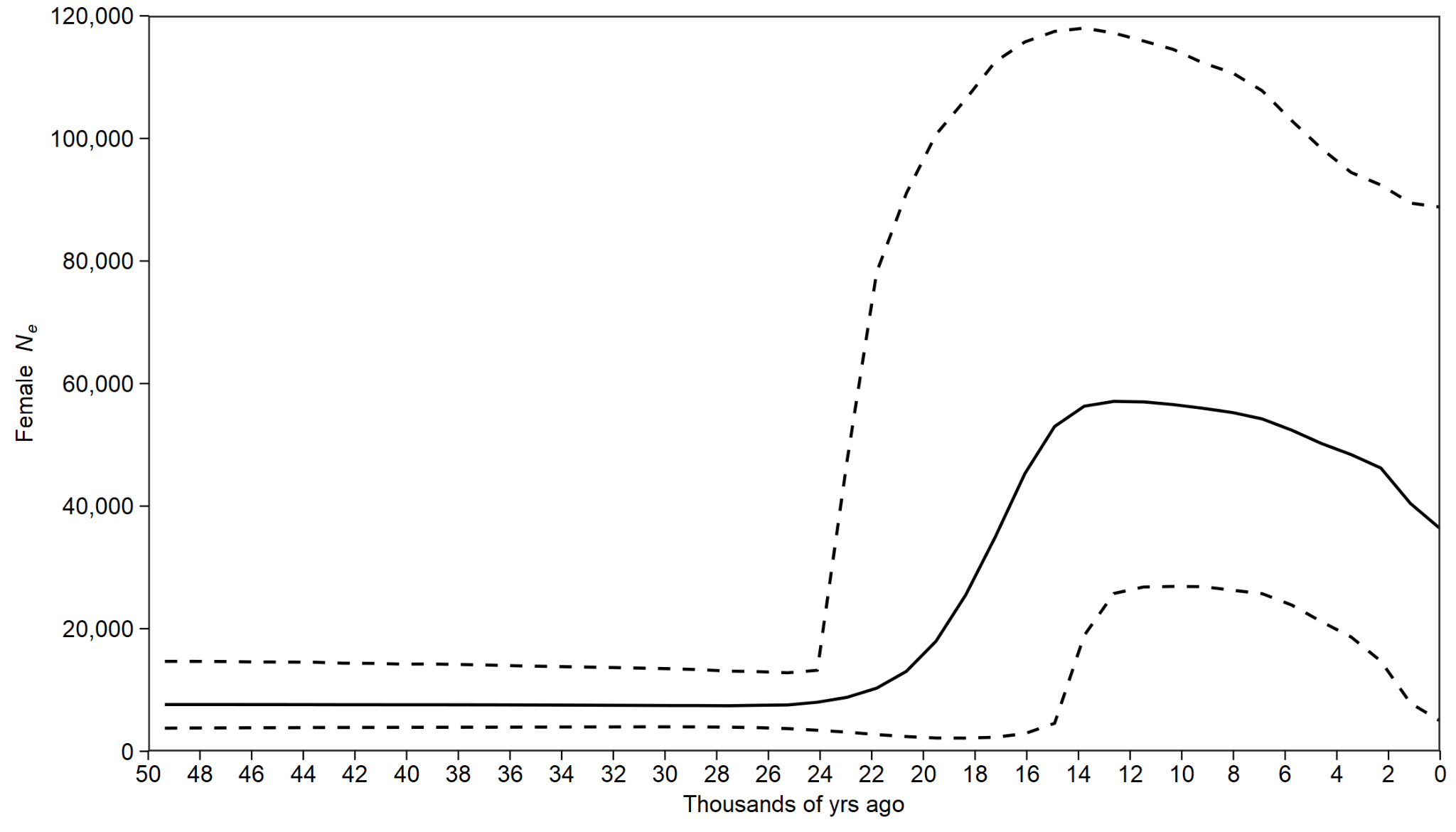

**Supplementary figure S20.** Bayesian Skyline plot for bowhead whales based on 129 complete mitochondrial genomes and calibrated using mean radiocarbon dates as tip dates in BEAST. Dotted lines show the 95% credibility intervals and the full line shows the mean value. An overview of the ages of the samples for which mitogenomes were retrieved is provided in Supplementary figure S19.

**Supplementary tables**

**Supplementary table S1.** Levels of heterozygosity and the significance of differences in mean values between groupings. Values given NA represent pairwise comparisons against themselves so no calculation was conducted. ‘Corrected fossil’ shows the results when adding a correction factor of -0.008, based on the biases observed with simulated data. Values in the Mann-Whitney-Wilcoxon Test below the greyed-out NA show the p-values

|  |  | **Fossils** | **Contemporary Svalbard** | **Contemporary Canada** | **Corrected fossils** |
| --- | --- | --- | --- | --- | --- |
|  | **Sample size** | 44 | 12 | 7 | 44 |
|  | **Mean** | 0.467 | 0.450 | 0.460 | 0.459 |
|  | **SD** | 0.0062 | 0.0082 | 0.0083 | 0.0062 |
| Mann-Whitney-Wilcoxon Test results | **Fossils** | NA |  |  |  |
|  | **Contemporary Svalbard** | 2.20E-16 | NA |  |  |
|  | **Contemporary Canada** | 2.20E-16 | 2.20E-16 | NA |  |
|  | **Corrected fossils** | NA | 2.20E-16 | 0.02014 | NA |

**Supplementary table S2.** Nuclear genome-wide nucleotide diversity, calculated by pooling individuals into windows of 1,000 year time spans. Calculated in 500 kb sliding windows. For the pre-whaling fossil samples, localities were pooled due to lacking genetic differentiation between Canada and Svalbard. Individuals with radiocarbon dates that could not be accurately calibrated due to the marine-reservoir effect were added to the 0-1,000 bin. Values are also shown visually in Supplementary figure S1.

| **Sample age (years BP)** | **Sample size** | **Mean pi** | **SD** |
| --- | --- | --- | --- |
| Contemporary Canada | 7 | 0.366 | 0.042 |
| Contemporary Svalbard | 12 | 0.355 | 0.035 |
| 0-1,000 | 7 | 0.365 | 0.037 |
| 1,000-2,000 | 5 | 0.364 | 0.045 |
| 2,000-3,000 | 4 | 0.364 | 0.046 |
| 3,000-4,000 | 3 | 0.362 | 0.057 |
| 4,000-5,000 | 3 | 0.365 | 0.059 |
| 5,000-6,000 | 7 | 0.366 | 0.038 |
| 6,000-7,000 | 1 | NA | NA |
| 7,000-8,000 | 1 | NA | NA |
| 8,000-9,000 | 4 | 0.365 | 0.049 |
| 9,000-10,000 | 7 | 0.365 | 0.038 |
| 10,000-11,000 | 2 | 0.362 | 0.109 |

**Supplementary table S3**: Significance of differences between pairwise comparisons of nuclear genome-wide diversity estimates calculated using a Mann-Whitney-Wilcoxon Test in R v4.1.1 ^9^. We used a Bonferroni correction to identify the threshold for significance (p-value of 0.05/6038 windows), giving us an upper p-value for significance of 0.000008. Comparisons with p-values <0.000008 are indicated in green.

**Supplementary table S4.** Mitochondrial haplotype and nucleotide diversity calculated in 1,000 year time bins. For the pre-whaling fossil samples, localities were pooled due to lacking genetic differentiation between Canada and Svalbard. Values are shown visually in Supplementary figure S1.

| **Sample age (years BP)** | **n** | **Haplotype diversity** | **Haplotype diversity SD** | **Nucleotide diversity** | **Nucleotide diversity SD** |
| --- | --- | --- | --- | --- | --- |
| Contemporary Canada | 7 | 1 | 0.076 | 0.0027 | 0.0005 |
| Contemporary Svalbard | 12 | 0.993 | 0.040 | 0.00292 | 0.00017 |
| 0-1,000 | 20 | 1 | 0.016 | 0.00243 | 0.00024 |
| 1,000-2,000 | 9 | 1 | 0.052 | 0.00191 | 0.00054 |
| 2,000-3,000 | 10 | 1 | 0.002 | 0.00218 | 0.00051 |
| 3,000-4,000 | 7 | 1 | 0.076 | 0.00282 | 0.00054 |
| 4,000-5,000 | 7 | 1 | 0.076 | 0.00192 | 0.00034 |
| 5,000-6,000 | 10 | 1 | 0.045 | 0.00298 | 0.00030 |
| 6,000-7,000 | 4 | 1 | 0.177 | 0.00239 | 0.00053 |
| 7,000-8,000 | 3 | 1 | 0.0272 | 0.00328 | 0.00098 |
| 8,000-9,000 | 10 | 1 | 0.045 | 0.00269 | 0.00027 |
| 9,000-10,000 | 14 | 1 | 0.027 | 0.00236 | 0.00045 |
| 10,000-11,000 | 7 | 1 | 0.076 | 0.00254 | 0.00046 |
| 11,000-12,000 | 2 | 1 | 0.5 | 0.00429 | 0.00214 |

**Supplementary table S5.** List of the 18 sites found across 16 genes showing the highest likelihoods for allele frequency change correlating with time after filtering for those found within protein coding genes.

| **Scaffold** | **Site** | **Putative gene name** | **Bowhead annotation name** |
| --- | --- | --- | --- |
| scaffold_278 | 23961 | MALT1 | bmy_07070 |
| scaffold_245 | 1507931 | SELENOW | bmy_06613 |
| scaffold_2521 | 228275 | SHCBP1 | bmy_20555 |
| scaffold_1846 | 28679 | FAM47E | bmy_18610 |
| scaffold_357 | 77143 | SLC35D3 | bmy_08151 |
| scaffold_2309 | 235002 | FLCN | bmy_20079 |
| scaffold_1452 | 407215 | TPRA1 | bmy_17084 |
| scaffold_2500 | 110012 | C5H4orf33 | bmy_20470 |
| scaffold_215 | 1603004 | FRAS1 | bmy_05828 |
| scaffold_42 | 1335296 | ADAM23 | bmy_01676 |
| scaffold_521 | 342226 | KCNT2 | bmy_10115 |
| scaffold_136 | 1655180 | PDE1A | bmy_04301 |
| scaffold_424 | 270137 | CPOX | bmy_08966 |
| scaffold_424 | 270139 | CPOX | bmy_08967 |
| scaffold_1315 | 315029 | SMOX | bmy_16224 |
| scaffold_95 | 1432243 | GPATCH4 | bmy_03422 |
| scaffold_1320 | 504332 | XRCC5 | bmy_16453 |
| scaffold_1320 | 506797 | XRCC5 | bmy_16454 |

**Supplementary table S6.** Putative phenotypes associated with the genes found to have sites with allele changes highly correlated with time. GWAS (Genome-wide association study) and MGI (mouse genome informatics) phenotypes taken from genecards.org. The table is attached as a spreadsheet.

**Supplementary table S7:** the mean +/- 1SD for sea surface temperature (SST) and sea ice cover (SIC), under ssp2-4.5 for the Canadian and Svalbard localities at the years 2050, 2070, and 2090.

| Region | Year | Variable | Scenario | Mean | SD |
| --- | --- | --- | --- | --- | --- |
| Canada | 2050 | SST | ssp2-4.5 | 2.1 | 3.2 |
| Canada | 2070 | SST | ssp2-4.5 | 2.6 | 3.6 |
| Canada | 2090 | SST | ssp2-4.5 | 3.1 | 3.9 |
| Svalbard | 2050 | SST | ssp2-4.5 | 4.2 | 2.5 |
| Svalbard | 2070 | SST | ssp2-4.5 | 5.8 | 2.7 |
| Svalbard | 2090 | SST | ssp2-4.5 | 6.8 | 3.9 |
| Svalbard | 2050 | SIC | ssp2-4.5 | 0.2 | 0.2 |
| Svalbard | 2070 | SIC | ssp2-4.5 | 0.1 | 0.2 |
| Svalbard | 2090 | SIC | ssp2-4.5 | 0 | 0.1 |
| Canada | 2050 | SIC | ssp2-4.5 | 0.3 | 0.3 |
| Canada | 2070 | SIC | ssp2-4.5 | 0.3 | 0.3 |
| Canada | 2090 | SIC | ssp2-4.5 | 0.3 | 0.3 |

**Supplementary table S8:** Mean +/- 1SD for sea surface temperature (SST) and sea ice cover (SIC), under ssp5-8.5 for the Canadian and Svalbard localities for the years 2050, 2070, and 2090.

| Region | Year | Variable | Scenario | Mean (^o^C) | SD |
| --- | --- | --- | --- | --- | --- |
| Canada | 2050 | SST | ssp5-8.5 | 2.4 | 3.4 |
| Canada | 2070 | SST | ssp5-8.5 | 3.4 | 4.1 |
| Canada | 2090 | SST | ssp5-8.5 | 5.1 | 4.8 |
| Svalbard | 2050 | SST | ssp5-8.5 | 4.6 | 2.6 |
| Svalbard | 2070 | SST | ssp5-8.5 | 7 | 2.9 |
| Svalbard | 2090 | SST | ssp5-8.5 | 9.4 | 3.2 |
| Canada | 2050 | SIC | ssp5-8.5 | 0.3 | 0.3 |
| Canada | 2070 | SIC | ssp5-8.5 | 0.2 | 0.3 |
| Canada | 2090 | SIC | ssp5-8.5 | 0.1 | 0.2 |
| Svalbard | 2050 | SIC | ssp5-8.5 | 0.1 | 0.1 |
| Svalbard | 2070 | SIC | ssp5-8.5 | 0 | 0.1 |
| Svalbard | 2090 | SIC | ssp5-8.5 | 0 | 0 |

**Supplementary table S9:** Radiocarbon ID, ages, coordinates, and references of the fossils used to inform the habitat modelling. The table includes 151 radiocarbon dates new to this study. yBP - years before present - attached as spreadsheet

**Supplementary table S10:** Bone collagen stable *δ*^13^C and *δ*^15^N isotope values for 200 pre-whaling fossil specimens. Data from 196 Holocene specimens were used in our analysis. Four individuals were Late Pleistocene, and were not included in our analysis. Attached as spreadsheet

**Supplementary table S11:** Sample information and mapping results for the initial genomic test shotgun sequencing of our pre-whaling Holocene and Late Pleistocene specimens - attached as spreadsheet.

**Supplementary table S12:** Mapping results of the 56 pre-whaling Holocene individuals that underwent further shotgun sequencing after the original test sequencing - attached as spreadsheet.

**Supplementary table S13:** Sample information and mapping results of the contemporary individuals, which included 7 from Canada and 12 from Svalbard - attached as spreadsheet

**Supplementary table S14.** SNP heterozygosity estimates of three high-coverage Svalbard individuals with downsampling and simulated ancient DNA damage patterns. ‘All sites’ is the heterozygosity estimated when using all sites at once. ‘Mean (20x500k)’ is the mean value of heterozygosity taken from 500k sites randomly sampled 20 times. ‘SD (20x500k)’ is the standard deviation of heterozygosity taken from 500k sites randomly sampled 20 times. Data is also presented visually in Supplementary figure S11.

| **Individual** | **Value** | **Original** | **2x** | **aDNA** | **aDNA 1x** | **aDNA 0.2x** |
| --- | --- | --- | --- | --- | --- | --- |
| 17-07 | All sites | 0.453 | 0.452 | 0.463 | 0.458 | 0.460 |
|  | Mean (20x500k) | 0.453 | 0.452 | 0.463 | 0.459 | 0.460 |
|  | SD (20x500k) | 0.001 | 0.002 | 0.001 | 0.003 | 0.011 |
| 17-12 | All sites | 0.448 | 0.450 | 0.459 | 0.460 | 0.458 |
|  | Mean (20x500kb) | 0.448 | 0.450 | 0.459 | 0.460 | 0.458 |
|  | SD (20x500kb) | 0.002 | 0.002 | 0.001 | 0.002 | 0.010 |
| 17-19 | All sites | 0.451 | 0.453 | 0.459 | 0.459 | 0.449 |
|  | Mean (20x500kb) | 0.451 | 0.452 | 0.459 | 0.459 | 0.452 |
|  | SD (20x500kb) | 0.001 | 0.002 | 0.002 | 0.003 | 0.008 |

**Supplementary table S15.** Alleles found in the contemporary bowhead individuals at the sites with allele changes highly correlated with time in genes putatively related to body height. ‘Al.’ means allele.

|  |  |  |  |  | **Svalbard (n=12)** | | **Canada (n=7)** | |
| --- | --- | --- | --- | --- | --- | --- | --- | --- |
| **Scaffold** | **Site** | **Gene** | **Allele 1** | **Allele 2** | **n Al. 1** | **n Al. 2** | **n Al. 1** | **n Al. 2** |
| scaffold_278 | 23961 | MALT1 | C | A | 10 | 2 | 1 | 1 |
| scaffold_245 | 1507931 | SELENOW | G | A | 8 | 4 | 1 | 1 |
| scaffold_2521 | 228275 | SHCBP1 | T | C | 6 | 6 | 1 | 1 |
| scaffold_1846 | 28679 | FAM47E | C | T | 5 | 7 | 2 | 2 |
| scaffold_357 | 77143 | SLC35D3 | G | A | 3 | 9 | 0 | 1 |
| scaffold_2309 | 235002 | FLCN | A | G | 5 | 7 | 3 | 2 |
| scaffold_1452 | 407215 | TPRA1 | T | C | 5 | 7 | 2 | 2 |
| scaffold_2500 | 110012 | C5H4orf33 | T | A | 6 | 6 | 1 | 3 |
| scaffold_215 | 1603004 | FRAS1 | T | G | 7 | 5 | 2 | 1 |
| scaffold_42 | 1335296 | ADAM23 | C | T | 7 | 5 | 0 | 4 |
| scaffold_521 | 342226 | KCNT2 | A | G | 5 | 7 | 1 | 2 |
| scaffold_136 | 1655180 | PDE1A | G | A | 10 | 2 | 1 | 1 |
| scaffold_424 | 270137 | CPOX | A | G | 9 | 3 | 4 | 0 |
| scaffold_424 | 270139 | CPOX | A | G | 9 | 3 | 4 | 0 |
| scaffold_1315 | 315029 | SMOX | C | T | 2 | 10 | 0 | 2 |
| scaffold_95 | 1432243 | GPATCH4 | G | C | 4 | 8 | 1 | 1 |
| scaffold_1320 | 504332 | XRCC5 | A | G | 5 | 7 | 0 | 2 |
| scaffold_1320 | 506797 | XRCC5 | G | A | 6 | 6 | 0 | 0 |

**Supplementary table S16:** Mapping results of the pre-whaling Holocene fossil individuals from Svalbard that underwent mitochondrial enrichment - attached as spreadsheet

**Supplementary video S1:** Reconstruction of spatiotemporal habitat suitability for bowhead whales over the last 11,000 years. Animation shows habitat suitability for the bowhead whale as predicted by our MaxEnt model and sea surface temperatures for the two focal regions in this study.
